# Characterising the metabolic rewiring of extremely slow growing *Komagataella phaffii*

**DOI:** 10.1101/2023.09.04.556224

**Authors:** Benjamin Luke Coltman, Corinna Rebnegger, Brigitte Gasser, Jürgen Zanghellini

**Affiliations:** CD-Laboratory for Growth-decoupled, Protein Production in Yeast at Department of Biotechnology, University of Natural, Resources and Life Sciences (BOKU), Vienna, Austria; University of Natural Resources and Life, Sciences (BOKU), Department of Biotechnology, Institute of Microbiology, and Microbial Biotechnology, Vienna, Austria; Austrian Centre of Industrial, Biotechnology, Vienna, Austria; Department of Analytical Chemistry, University of Vienna, Vienna, Austria

**Keywords:** Metabolism, Modelling, *K. phaffii*, Near-zero growth, FBA, Flux Sampling

## Abstract

Retentostat cultivations have enabled investigations into substrate-limited near-zero growth for a number of microbes. Quantitative physiology at these near-zero growth conditions has been widely discussed, yet characterisation of the fluxome is relatively under-reported. We investigated the rewiring of metabolism in the transition of a recombinant protein producing strain of *Komagataella phaffii* to glucose-limited near-zero growth rates. We used cultivation data from a 200-fold range of growth rates and comprehensive biomass composition data to integrate growth rate dependent biomass equations, generated using a number of different approaches, into a *K. phaffii* genome scale metabolic model. Here we show that a non-growth associated maintenance value of 0.65 mmol_ATP_ g_CDW_^−1^ h^−1^ and a growth-associated maintenance value of 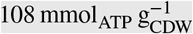 lead to accurate growth-rate predictions. In-line with its role as energy source, metabolism is rewired to increase the yield of ATP per glucose. This includes a reduction of flux through the pentose phosphate pathway, and a greater utilisation of glycolysis and the TCA cycle. Interestingly, we observed activity of an external, non-proton translocating NADH dehydrogenase in addition to the malate-aspartate shuttle. Regardless of the method used for the generation of biomass equations, a similar, yet different, growth rate dependent rewiring was predicted. As expected, these differences between the different methods were clearer at higher growth rates, where the biomass equation provides a much greater constraint than at slower growth rates. When placed on an increasingly limited glucose diet, the metabolism of *K. phaffii* adapts, enabling it to continue to drive critical processes sustaining its high viability at near-zero growth rates.

## INTRODUCTION

Microorganisms are regularly exposed to variable nutrient availability, both in their natural environments and when employed in bioprocesses. Understanding the physiological responses of microorganisms to these conditions aids our understanding of life in extreme environments and improves the efficiency of bioprocess design. Microbial hosts frequently exhibit a dependency of (heterologous) product formation rates on cell proliferation, which currently requires a bioprocess to be operated at high growth rates to achieve high titers (Z. Liu, Hou, Martínez, Petranovic, & Nielsen 2013; Looser et al. 2015). However, at higher cell densities this becomes infeasible. Subsequently, a typical bioprocess achieves high titers via high-density cultivations at slower growth rates (Buchetics et al. 2011; Maurer, Kühleitner, Gasser, & Mattanovich 2006). An ideal bioprocess would minimise loss of substrate to unwanted by-products. For secreted products, this includes biomass generation. Continuous cultivation at very low or near-zero growth rates can potentially uncouple these complex traits, increasing product yields and avoiding losses to biomass.

*Komagataella phaffii* (syn. *Pichia pastoris*), a methylotrophic Crabtree-negative budding yeast, is a well-adopted host for recombinant protein production due to the typical benefits of a eukaryotic expression host, reasonably efficient protein production rates and its ability to be cultivated at very high cell densities (Cregg, Cereghino, Shi, & Higgins 2000; Gasser et al. 2013; Heyland, Fu, Blank, & Schmid 2010; W.-C. Liu et al. 2019; Mattanovich, Sauer, & Gasser 2017). While *K. phaffii* can utilise methanol as the sole carbon and energy source, methanol-based production also comes with disadvantages, thus glucose is often the preferred nutrient in bioprocesses (Wang et al. 2017; Yang & Zhang 2018). Physiological changes have been investigated across a range of different growth rates in *K. phaffii* (Garcia-Ortega, Adelantado, Ferrer, Montesinos, & Valero 2016; Heyland et al. 2010; Rebnegger et al. 2014), and were further exploited for bioprocess design (Maurer et al. 2006). Yet very slow growth rates (below 0.025 h^−1^) have been underrepresented in these studies. In principle, very slow growth rates can be achieved in fed-batch cultivations, although their applicability for studying cell physiology is limited. Operating chemostats at very slow growth rates becomes infeasible, whilst retentostat cultivations enable the study of organisms at extremely slow growth rates. Retentostats are modified chemostats, where complete biomass retention is achieved through effluent outflow through a filter. The combination of this biomass retention and a constant substrate feed enables the transition to very slow growth rates. If the limiting substrate is the energy substrate, the biomass-specific substrate uptake rate *q*_s_ steadily decreases until it is equal to the biomass-specific energy substrate requirement for maintenance *m*_s_, as represented by the Pirt equation (Pirt 1982),

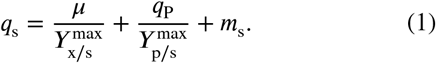

This substrate limitation is inherently different to a sudden substrate depletion, ensuring cells are not starving and remain both viable and metabolically active (L. G. Boender et al. 2011; L. G. M. Boender, de Hulster, van Maris, Daran-Lapujade, & Pronk 2009).

A number of physiological assessments have been made from retentostat cultivations for different yeast species, including: cell morphology, biomass compositions, and, transcriptome (L. G. Boender et al. 2011; Juergens et al. 2020; Rebnegger et al. 2016; Vos et al. 2016) and proteome analysis (Binai et al. 2014; Juergens et al. 2020). Glucose-limited retentostat cultivations of the Crabtree-positive yeast *Saccharomyces cerevisiae* have been performed in both aerobic (Bisschops et al. 2017; Vos et al. 2016) and anaerobic (Binai et al. 2014; L. G. Boender et al. 2011; L. G. M. Boender et al. 2009) conditions. Furthermore, non-energy-limited cultivations with nitrogen and phosphorous source limitation have been performed (Y. Liu, el Masoudi, Pronk, & van Gulik 2019). In ammonium-limited retentostat cultivations, production of succinic acid was uncoupled from growth, achieving a succinic acid yield on glucose of 0.61 mol mol^−1^ for over 500 h, albeit with accumulation of large quantities of inviable biomass (Y. Liu, Esen, Pronk, & van Gulik 2021). Aerobic glucose-limited retentostat cultivations were also performed to investigate two different Crabtree-negative methylotrophic yeasts, namely *K. phaffii* and *Ogataea polymorpha*, the latter being also thermotolerant. In comparison to *S. cerevisiae*, both retained a higher viability over a period of several weeks and both had a lower but growth rate dependent maintenance requirement (Juergens et al. 2020; Rebnegger et al. 2016; Vos et al. 2016).

The fluxome at very slow growth rates remains relatively under-reported, with a possible explanation being the complexities of adapting labelling experiments to retentostat cultivations. Constraint-based modeling (CBM) methods have been utilised for metabolic flux calculations from retentostat cultivations of the Gram-positive bacteria *Lactobacillus plantarum*, and more recently for non-energy substrate-limited retentostat cultures of *S. cerevisiae* (Goffin et al. 2010; Y. Liu et al. 2019). To employ CBM methods to estimate metabolic fluxes, an accurate genome-scale metabolic model (GSMM) constrained with exchange rates and a well-determined biomass composition are required (Dikicioglu, Kirdar, & Oliver 2015). For *K. phaffii*, a manually curated GSMM incorporated a consensus glucose biomass equation that reliably predicted growth rates at 0.1 h^−1^ under different oxygen limitations (Carnicer et al. 2009; Tomàs-Gamisans, Ferrer, & Albiol 2016). A later revision, *i*MT1026v3, added context-dependent biomass compositions for growth on methanol and glycerol, extending the model’s predictive abilities (Tomàs-Gamisans, Ferrer, & Albiol 2018).

Until recently, it was presumed that recombinant protein secretion would be almost infeasible in retentostat cultivation due to down-regulation of the secretory machinery at slow growth, a budding-associated secretion mechanism, and the lack of suitable promoters Puxbaum, Gasser, and Mattanovich (2016); Rebnegger et al. (2016); Wanka et al. (2016). However, Rebnegger et al. (2023) recently found that *K. phaffii* was actively secreting a recombinant bivalent nanobody (vHH) when expressed under control of the glucose-limit induced promoter P_G1-3_ in glucose-limited retentostat cultivations down to growth rates of *μ* = 0.00046 h^−1^. Whilst severe carbon and energy source limitation is not optimal for a bioprocess, we wanted to understand how the fluxome of the recombinant strain is affected at these extremely low growth rates. We expanded on a previous regression model of glucose-limited retentostat cultivations and used biomass composition data to generate multiple growth rate-specific biomass equations, using 3 different methods. Accurate metabolic model growth rate predictions were obtained by fitting the Pirt equation to maximal ATP production rates throughout the retentostat cultivation. Generated flux distributions demonstrated that the rewiring of metabolism could largely be attributed to increasing the yield of ATP per glucose metabolised. We also observed differences in flux distributions dependent on which method was used to generate the biomass equations, although the general trends in rewiring were similar. Our findings contribute to understanding the very low, growth rate dependent *m*_s_ observed in *K. phaffii*.

## RESULTS

### Incorporating a dynamic death rate into the retentostat regression model

Accurate exchange rates were generated using cultivation data from Rebnegger et al. (2023) and an extension of the regression model (Rebnegger et al. 2016). Throughout the retentostat cultivation, Rebnegger et al. (2023) measured the: (i) glucose concentration entering the retentostat *C*_S,in_; (ii) the total biomass concentration *C*_X_; (iii) the culture viability v; and, (iv) the concentration of the secreted recombinant protein vHH *C*_P,R_. We used this data (initial 0.025 h^−1^ chemostat and 10 sample points) to fit a regression model describing the retentostat cultivation. The model was based on a previous representation, but expanded to incorporate a dynamic death rate and account for protein production without the use of a predetermined relationship between biomass-specific productivity *q*_p_ and growth rate *μ* (Rebnegger et al. 2023 2016).

The retentostat cultivation is described by a system of ordinary differential equations (ODEs) describing the changing concentrations for viable biomass

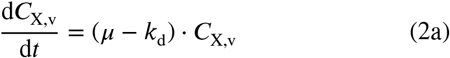

inviable biomass

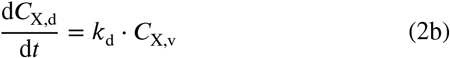

and product of interest

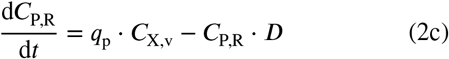

where *D* and *k*_d_ represent the dilution rate and death rate, respectively. *μ* and *q*_p_ are not independent of each other but connected via the Pirt equation (Pirt 1982),

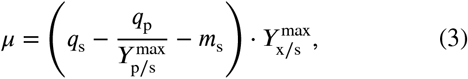

where *m*_s_, *q*_s_, 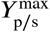, and 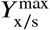 denote the substrate-specific maintenance coefficient, biomass-specific substrate uptake rate, maximum product yield and maximum biomass yield, respectively. *C*_S,in_ is controlled via a mixing vessel fed by two media reservoirs, *C*_S,MC_ and *C*_S,MR_ in either the chemostat or retentostat phase, respectively. During the retentostat, *C*_S,in_ approaches *C*_S,MR_ according to

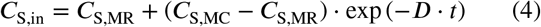

As glucose is limiting, the concentration of the glucose feed (*C*_S,in_) is much greater than the concentration in the retentostat (*C*_S,R_), *C*_S,in_ ≫ *C*_S,R_ and we assume a pseudo steady state 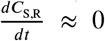 . Residual glucose concentrations were extremely low, *C*_S,R_ ≈ 0 (Rebnegger et al. 2023). *q*_s_ is therefore given by:

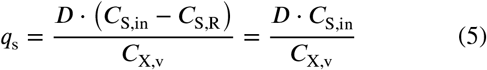

To compute

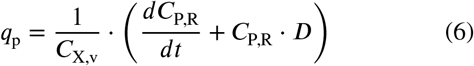

for each cultivation, the experimentally observed *C*_P,R_ = *p*_4_(*t*; 4) was fitted with a polynomial *p*_4_ of order 4 as a function of time, *t*. The function was incorporated into the series of ODEs to evaluate *q*_p_, during the cultivation.

Similarly, to compute *k*, the measured culture viability v as used,

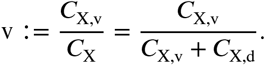

Using (2) and the quotient rule, it was found that

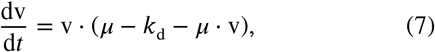

which leads to

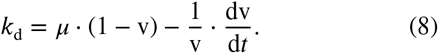

For each separate cultivation, the experimentally observed v = *p*_3_(*t*; 3) was fitted with a polynomial *p*_3_ of order 3 as a function of time, *t*. This function was incorporated into the series of ODEs to evaluate *k*_d_ during the cultivation. In situations where (8) was negative, the value was set to 0, preventing “resurrection” of dead biomass. Parameters and settings used during fitting are summarised in Table S1.

*m*_s_ was separately estimated for each cultivation through weighted least-squares regression between the predicted and experimental values of *C*_X_ and *C*_X,v_. The average estimated *m*_s_ between the three retentostat cultivations was 3.65 ± 0.11 mg g^−1^ h^−1^. Total *R*^2^, taking into account both *C*_X_ and *C*_X,v_ remained high for both models (> 0.999), whereas the *R*^2^ of the dead biomass *C*_X,d_ improved after conversion of *k*_d_ to a dynamic parameter (> 0.9 for the dynamic model compared to 0.45 to 0.99 for the model with a static death rate, Table S2). Both *q*_s_ and *q*_P_ were derived from the regression model, whilst the integrated biomass concentrations were used for calculating biomass-specific gas exchange rates from mass balancing. There was a lack of concordance in the oxygen exchange rates calculated from the separate reactors in both chemostat and retentostat phase, subsequently, at each sampling point we assumed a relative error of 50 % in 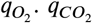 was in better agreement between replicates, however, below 0.0022 h^−1^ the initial flux balance analysis (FBA) solutions were infeasible without either secreting a carbon-containing compound or accepting an error on the carbon dioxide production rate. As the OUR and CPR are low at slow growth rates and no secreted compounds were detected in HPLC, we assumed that the gas detection system was not sensitive enough for the low gas exchange rates and subsequently assumed an error of 50 % for the *q*_*CO*_ below a growth rate of 0.0022 h^−1^.

### Generation of growth rate dependent biomass equations

To use CBM methods, we generated growth rate specific stoichiometric representations of biomass formation, the so called biomass equation, of the recombinant protein producing *K. phaffii* strain.

Rebnegger et al. (2023) analysed the biomass composition (from 3 biological replicates) at 5 different growth rates: a chemostat cultivation at 0.1 h^−1^, the chemostats preceding the retentostat cultivation at 0.025 h^−1^ and three sampling points during the retentostat cultivation (see black arrows in central panel of Figure 1). Glycogen and trehalose content were also measured at 7 additional sampling points during the retentostat cultivation, where additional process parameters were also determined: *C*_X_, *C*_P,R_, v and *C*_S,in_ (depicted by circles in Figure 1) (Table S3). The biomass composition is typically analysed from steady-state cultures, and is often combined with assumptions from literature to generate a GSMM biomass equation (Carnicer et al. 2009; Széliová et al. 2020; Tomàs-Gamisans et al. 2018). For all 5 analysed biomass compositions, we combined them with assumptions previously used in the generation of the consensus glucose biomass equation in *i*MT1026v3 (see Supplementary Methods S1.2, Table S4) (Tomàs-Gamisans et al. 2016). To generate the growth rate specific biomass equations for 0.1 h^−1^ and 0.025 h^−1^, the combined compositions were used directly as the biomass equation.

**FIGURE 1.**
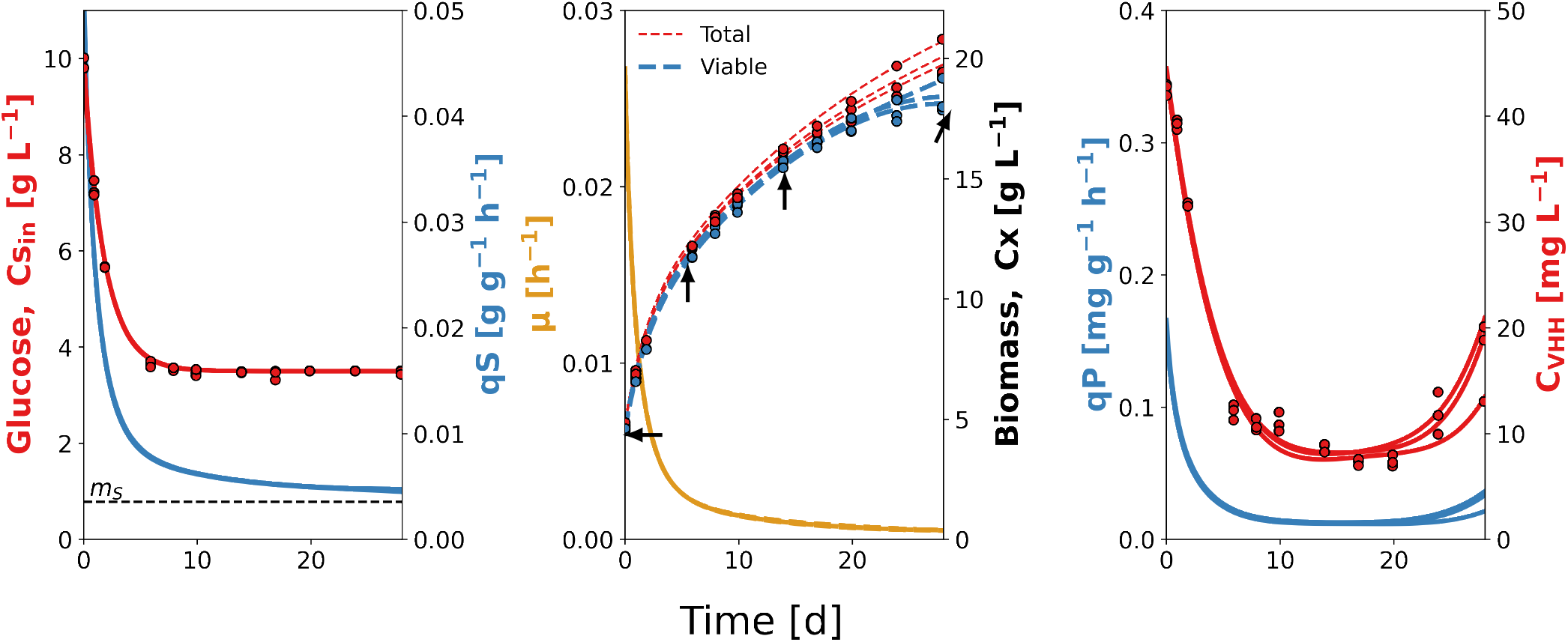
Accumulation profiles and derivative rates of the three retentostat cultivations after regression model adaptations. Left: Glucose concentration of feed media (*C*_S,in_, red), biomass-specific glucose uptake rates (*q*_s_, blue) and the average specific maintenance coefficient (*m*_s_ = 3.56 mg g^−1^ h^−1^, black dotted line). Centre: Total (*C*_X_, red) and viable (*C*_X,v_, blue) biomass concentrations and the specific growth rate (*μ*, yellow). Black arrows highlight sampling points where biomass composition was measured. Right: Concentration of recombinant secreted protein (vHH) in the retentostat (*C*_P,R_, red) and the biomass-specific vHH production rate (*q*_P_, blue). Scatter points represent measured values.

As the retentostat cultivation is in pseudo-steady state, we evaluated 3 different methods to generate the 10 additional biomass equations (an equation for each sampling point during the retentostat cultivation). All 3 methods represent the 4 biomass compositions (initial chemostat and 3 later points during retentostat cultivation) against time since retentostat initiation, but differ in how they determine the changing composition and subsequently the stoichiometry of the biomass equation. We refer to these methods as: (i) derived, (ii) fitted, and (iii) interpolated.

The “interpolated” equations were generated by representing the compositions against time, and a linear interpolation for each component between the data points. Subsequently, the stoichiometry of both the generated equation and the biomass composition were equal at the 3 sampling points of the retentostat.

Both the “fitted” and “derived” equations were generated from representing the concentration of each biomass component in the reactor against time. First, the concentration of each biomass component, 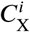, was calculated during the retentostat cultivation based on *C*_X_ from the regression model:

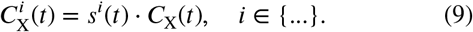

where *i* represents a component of biomass and *s*^*i*^ its stoichiometric coefficient. It was observed that, except for glycogen, the concentrations of all other biomass components increase monotonically with time, which was well captured by fitting an exponential asymptotic function (11) (coefficient of determination > 0.95) (Figure S1).

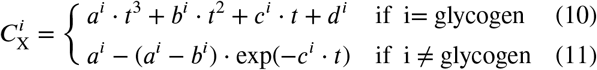

The glycogen concentration initially increased, before decreasing after approximately 200 h (Figure S1, see Table S3), and was better fitted with a cubic polynomial (adjusted coefficient of determination > 0.975) (Figure S1). The asymptotic function was chosen due to its monotonic characteristics which we deemed a reasonable assumption without further data. The asymptotic function provided a near linear fit for hexosylceramide and inositol-containing lipids. We assumed this was reasonable due to: 1) these lipids falling below the limit of quantification using external and internal standards; and, 2) the lack of lipidomics data for the sampling point at day ∼6, where an average between the day 0 and ∼14 was used.

The stoichiometric coefficients of the fitted equations were calculated via (12) using the fit (11) and (10).

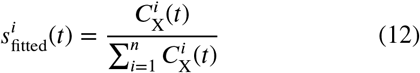

The derived equations were generated from the above fitted functions, using the assumption of Goffin et al. (2010), that the change in the concentration of biomass components is exclusively due to the generation of new biomass. Subsequently, 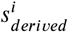 was calculated by solving (13) numerically, using the fit of (11) and (10).

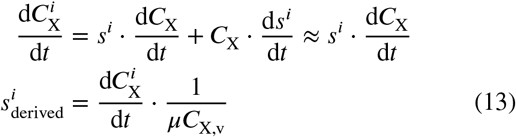

As the concentration of glycogen decreases at a certain point, this led to a negative stoichiometric coefficient in the derived equation, which was represented by an active sink reaction when using the CBM methods.

For each equation generated with the three methods, the oichiometric coefficients were scaled such that the biomass molecular weight was equal to 1 g mmol^−1^ (Chan, Cai, Wang, Simons-Senftle, & Maranas 2017; Dinh, Sarkar, & Maranas 2022). The regression model derived growth rates were used in the conversion of the sampling times to growth rate specific equations. As mentioned, the biomass component coefficients of the 0.1 h^−1^ and 0.025 h^−1^ biomass equations were the same for all three methods.

The calculated stoichiometries changed over the course of the cultivation and between the three generation methods, although the pattern of changes was similar. The equations generated with the interpolated and fitted methods were very similar, with the median relative stoichiometry between two growth rate specific equations varying between 96 % to 101 %. Relative to the interpolated and fitted equations, the derived equations varied more, between 83 % to 112 % and 83 % to 107 %, respectively. The stoichiometry of a number of components varied much more with the derived equations, particularly at slower growth rates (Figure S2 and Table S5). These components included alpha,alpha-trehalose, glutamate and glutamine, ergosteryl and zymosterol ester, triglycerides and ceramides. These differences were also evident on the macromolecular level (Figure S3). For all three methods, the carbohydrate and protein content were similar and stable across all growth rates; the lipid content decreased slightly before increasing at slower growth rates; and, both the DNA and RNA content reduced at slower growth rates. In comparison with the consensus biomass equation, the carbohydrate and DNA content were generally higher for all methods; RNA and lipid content generally lower and the protein content similar. As the consensus equation was determined at 0.1 h^−1^, a direct comparison with our 0.1 h^−1^ equation shows that the carbohydrate and DNA content of our equation were higher, RNA and lipid content lower and the protein content almost identical (Figure S3). An additional biomass equation, which we refer to as scaled consensus, was also generated, where the stoichiometric coefficients of the consensus biomass equation were scaled to sum to 1 g mmol^−1^. This led to a 15 % increase in the stoichiometry of all components. Although the biomass component coefficients of the 0.1 h^−1^ and 0.025 h^−1^ biomass equation are the same for all three methods, the energetic parameters could vary between the different methods. Subsequently, an equation for 0.1 h^−1^ and 0.025 h^−1^ was defined for each method. The 36 (12 growth rates, 3 methods) growth rate specific biomass equations were incorporated into the modified model *i*MT1026-NZ, in addition to the scaled consensus equation.

### Accurate growth rate predictions with static energetic parameters

ATP is the primary energy currency of a cell; assuming limitation, its distribution between maintenance and growth can be represented via the Pirt equation (Pirt 1982):

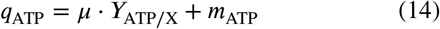

where *m*_ATP_ and *Y*_ATP/X_ denote non-growth associated maintenance (NGAM) and growth associated maintenance (GAM) demand, respectively. The latter represents the energy required for polymer synthesis and energy-depleting activities such as protein translocation and turnover, whereas NGAM accounts for processes such as maintenance of ion gradients across membranes and cell repair (Thiele & Palsson 2010). Here, we assume GAM additionally accounts for all non-polymerisation costs of recombinant protein production. Fitting these energetic parameters is important for prediction of both growth rates and intracellular fluxes (Széliová et al. 2021; Tomàs-Gamisans et al. 2016; Tomàs-Gamisans et al. 2018).

We predicted these parameters by fitting growth rate predictions of a constrained *i*MT1026-NZ to the retentostat cultivation data. We evaluated these parameters for each of the 3 biomass equation generation methods, in addition to using the previously published consensus glucose biomass equation and the scaled consensus equation (Tomàs-Gamisans et al. 2016).

At the 12 different growth rates, *i*MT1026-NZ was constrained with the average biomass-specific exchange fluxes, the biomass equation set to the respective equation and the model growth rate fixed to the average of the three regression model derived growth rates. Then, *q*_ATP_ was maximised via FBA (Figure 2a). A relative distance weighted linear function was fitted to the *μ*-*q*_ATP_ relationship, avoiding dominance of the significantly larger *q*_ATP_ at higher growth rates. The y-intercept, NGAM, was determined as 0.69 mmol_ATP_ g_CDW_^−1^ h^−1^ for the consensus equation, 0.65 mmol_ATP_ g_CDW_^−1^ h^−1^ for the derived equations, and 0.66 mmol_ATP_ g_CDW_^−1^ h^−1^ for the fitted, interpolated and scaled consensus equations.

**FIGURE 2.**
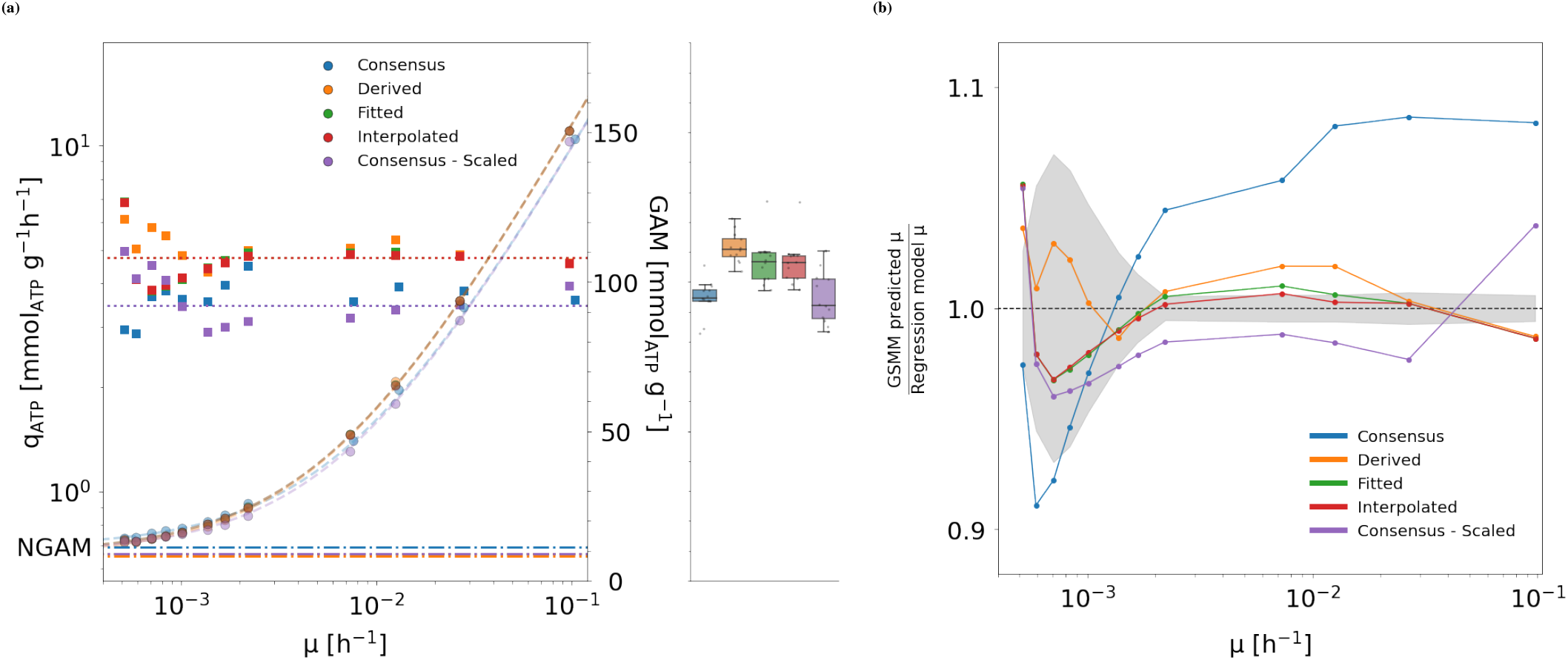
Energetic parameter fitting for each biomass equation generation method. a The *q*_ATP_ (circles) and growth associated maintenance (GAM) (squares) across all growth rates. *q*_ATP_ values are very similar between different methods and points often overlap. For each biomass generation method, a relative weighted linear function (dashed line) was fitted to the *q*_ATP_-*μ* relationship to determine the non-growth associated maintenance (NGAM) (lower dashed/dotted line) . At each growth rate, the GAM was calculated using the predicted *q*_ATP_ and the determined NGAM. The distribution of these GAM values is also represented in the box plot. A constant GAM (horizontal dotted line) was estimated from error weighted least-squares regression between regression model and *i*MT1026v3 predicted growth rates. b Comparison of the regression-model relative growth rate predictions for each type of biomass equation after determining optimal energetic parameters. The grey shaded area represents the standard deviation of the growth rate between the individual regression fits of the retentostat cultivation.

Next, we determined a growth rate independent GAM through fitting the *i*MT1026-NZ predicted growth rates to the regression model growth rates. An optimal GAM of 92 mmol_ATP_ g_CDW_^−1^ was determined for the consensus and scaled consensus biomass equation, and 108 mmol_ATP_ g_CDW_^−1^ for the derived, fitted and interpolated equations (Figure 2a). As the stoichiometry of the 0.1 h^−1^ and 0.025 h^−1^ biomass equations for the three different methods are the same, and the energetic parameters for the fitted and interpolated method are identical, then the 0.1 h^−1^ and 0.025 h^−1^ equations for these two methods are identical. An NGAM of ^∼^0.65 mmol_ATP_ g_CDW_^−1^ h^−1^ is calculated from the *m*_s_ of 3.65 mg g^−1^ h^−1^ using the maximum stoichiometric ATP-glucose yield of *i*MT1026-NZ (*Y*_ATP/S_ ≈ 32 mmol_ATP_ mmol_glucose_^−1^).

The growth rates predicted with all biomass equations were within 90 % to 110 % of the regression model predicted growth rates (Figure 2b). Growth rate predictions with the fitted, interpolated and derived equations were very similar to each other at all growth rates and were within 95 % to 105 % of the regression model. The scaled consensus equation more accurately predicted growth rates than the consensus equation, but less accurately than the other three methods. At 0.1 h^−1^, the consensus and scaled consensus equation over-predicted the growth rate. The sum of squared errors between the regression model growth rate and predictions with the consensus, scaled consensus, derived, fitted and interpolated equations were 694, 78, 29, 16 and 12, respectively. Thus, across the ∼200-fold range of investigated growth rates, the assumption of a growth rate independent GAM and NGAM of the Pirt equation led to reliable growth rate predictions. The GAM of each biomass equation in *i*MT1026-NZ was updated and the respective NGAM used in all further analysis.

### Metabolic rewiring during the transition to near-zero growth rates

Based on the reliability of the growth rate predictions, we investigated the changing flux distributions of the recombinant protein producing *K. phaffii* strain during the transition to glucose-limited near-zero growth. At each investigated growth rate, *i*MT1026-NZ was constrained with the relevant exchange rates and respective biomass equations before generating flux distributions. Single flux distributions were generated with pFBA, assuming growth maximisation as the objective; whilst the constrained solution space was characterised via flux sampling. pFBA generates a single solution, with minimal total absolute flux, reflecting optimal metabolic usage (Lewis et al. 2010). Flux sampling provides a less biased characterisation of the range of possible flux distributions as well as their probability, although ensuring sampler convergence is important (Supplementary Results S2.1) (Herrmann, Dyson, Vass, Johnson, & Schwartz 2019; Megchelenbrink, Huynen, & Marchiori 2014). When comparing the results between pFBA and flux sampling, ∼70 % of pFBA solutions fell within the 95 % confidence interval of the sampled fluxes. The results deviated between methods for reactions in a number of pathways, including: exchange reactions, amino sugar and nucleotide sugar metabolism, sterol metabolism and the TCA cycle. Across all growth rates and types of biomass equation, a median of ∼65 % and ∼78 % of pFBA solutions fell within the 95 % confidence interval for reactions in glycolysis and the pentose phosphate pathway (PPP), respectively. Yet, both pFBA and sampling revealed a similar growth rate dependent pattern of change.

To compare between different growth rates/glucose uptake rates, fluxes were normalised to the glucose uptake rate (flux yield) revealing a growth rate associated rewiring of central carbon metabolism (Figure 3 and Table S6). The flux predictions showed some biomass equation dependency, although the growth rate associated pattern of changes was shared between all biomass equations. Results with all three methods used for biomass equation generation were similar across all growth rates, but were different to those obtained with either the consensus equation or the scaled version. The largest changes occurred between 0.1 h^−1^ and 0.0022 h^−1^, representing the separate chemostat and the sampling point after ∼6 days of retentostat cultivation respectively; the absolute change in *q*_s_ was greatest in this period (Figure 1). Additionally, for some reactions we observed a large shift in the sampled fluxes between the two growth rates of 0.0073 h^−1^ and 0.0022 h^−1^. Between these two growth rates, the concentration of media entering the retentostat stabilises at ∼3.5 g L^−1^, representing the transition point of the substrate concentration of the feed media being exclusively defined by the retentostat media. A positive correlation in the flux yield was observed between glucose entry to the PPP via glucose-6-phosphate dehydrogenase (G6PDH2) and growth rate, and (consequently) a negative correlation between glycolysis entry and growth rate (Figure 3). At a growth rate of 0.00083 h^−1^, with the derived equation the median flux yield entering the PPP via G6PDH2 was ^∼^12 %, whilst glucose-6-phosphate isomerase (PGI) accounted for ^∼^81 %. At 0.1 h^−1^, 31 % entered PPP, and 44 % entered glycolysis via PGI (Figure 3). The pFBA results were similar (Figure 4).

**FIGURE 3.**
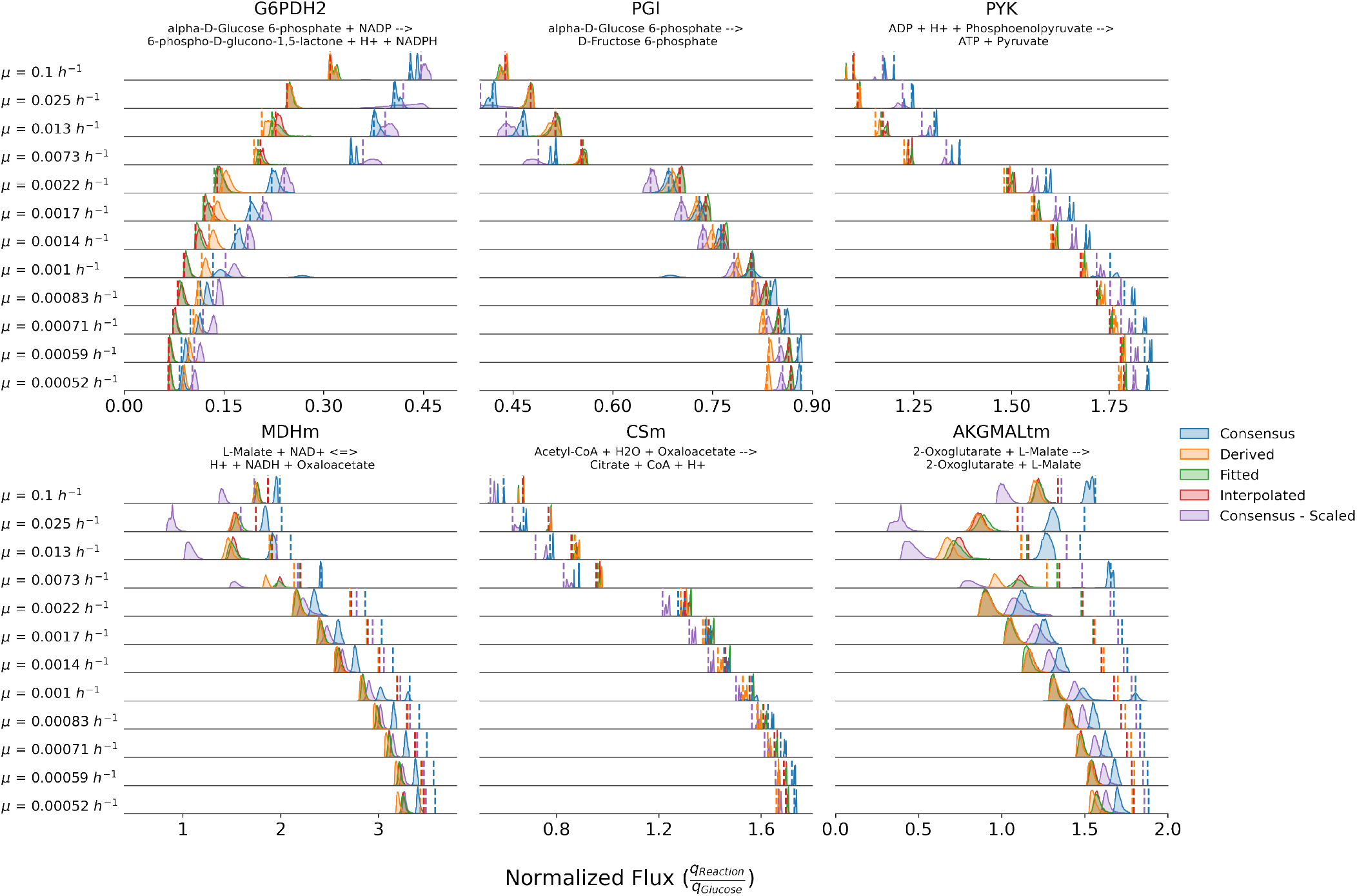
Glucose uptake normalised flux distributions for a number of branch points of central carbon metabolism. Dashed lines represent parsimonious flux balance analysis (pFBA) solutions, density plots are generated from flux sampling solutions. Different colours distinguish predictions generated using the different types of biomass equation and their respective energetic parameters.

**FIGURE 4.**
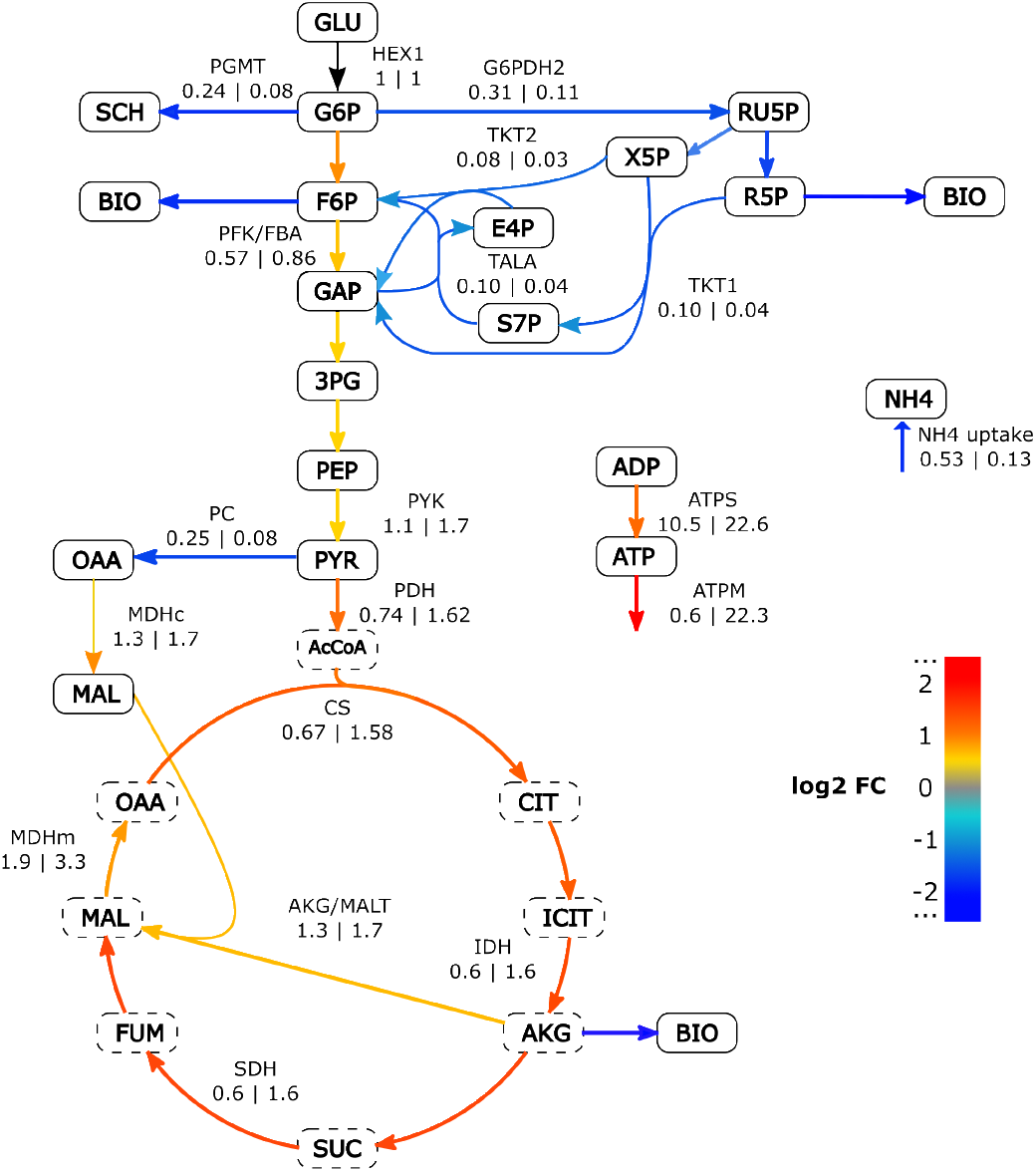
Map of central carbon metabolism comparing the metabolic flux distributions at 0.1 h^−1^ and 0.00083 h^−1^. Metabolic fluxes are normalised to glucose uptake flux (here represented as HEX1). At each growth rate, *i*MT1026-NZ was constrained with the relevant cultivation data and the growth rate specific derived biomass equation. The arrow color represents the log2 fold-change between predicted fluxes. Blue = reduced flux at the lower growth rate, red = increased flux at lower growth rate. At lower growth rates, less glucose is metabolised through the PPP, with more through glycolysis and the TCA cycle. Cytosolic and mitochondrial metabolites are represented with solid or dashed outlines, respectively. SCH = storage carbohydrates, BIO = branching point for biomass precursor synthesis.

In agreement with an increased glycolysis utilisation at slower growth rates, an increase in the flux yield of pyruvate kinase (PYK) and components of the TCA cycle was observed (Figure 3). There were changes in the flux yield of components of the malate aspartate shuttle. The flux yield of malate dehydrogenase (MDH) increased when cells approached near-zero growth, whereas the flux yield of the malate-alpha-ketoglutarate antiporter (AKGMALtm) initially reduced between 0.1 h^−1^ and 0.025 h^−1^, before increasing at slower growth rates. The shuttle regenerates reducing equivalents in the mitochondria, allowing electrons harvested during glycolysis to be passed to the electron transport chain.

Further changes were observed in the flux yield of many components of oxidative phosphorylation (Figure 5). The flux yields of respiratory complexes I (NADH2_u6mh), II (SUCD2_u6m), III (CYOR_u6m), IV (CYOOm) and ATP synthase (ATPS3m) increased at slower growth rates (Figure 5). The relative contribution of pathways to total ATP production changed slightly, with a small increase in the relative contribution of oxidative phosphorylation to ATP production during the transition to near-zero growth rates (Table S7). Between 0.1 h^−1^ and 0.025 h^−1^, the median flux yield of ATP synthase was similar ^∼^11 mmol_ATP_ mmol_glucose_^−1^. Yet, 4 times as much ATP was used to satisfy the NGAM (ATPM) at 0.025 h^−1^ than at 0.1 h^−1^ (Figure 5). At 0.00083 h^−1, ∼^37 times more ATP was used to satisfy the NGAM than at 0.1 h^−1^ (0.6 and 22.2 mmol_ATP_ mmol_glucose_^−1^, respectively), whilst the yield of ATP per glucose doubled (∼11 and 22.7 mmol_ATP_ mmol_glucose_^−1^, respectively)(Figure 4 and Figure 6). At the slowest analysed growth rates, the ATP synthase flux yield increased up to 25 mmol_ATP_ mmol_glucose_^−1^, but did not reach the stoichiometric maximum for *i*MT1026v3 of 32 mmol_ATP_ mmol_glucose_^−1^ (Tomàs-Gamisans et al. 2016).

**FIGURE 5.**
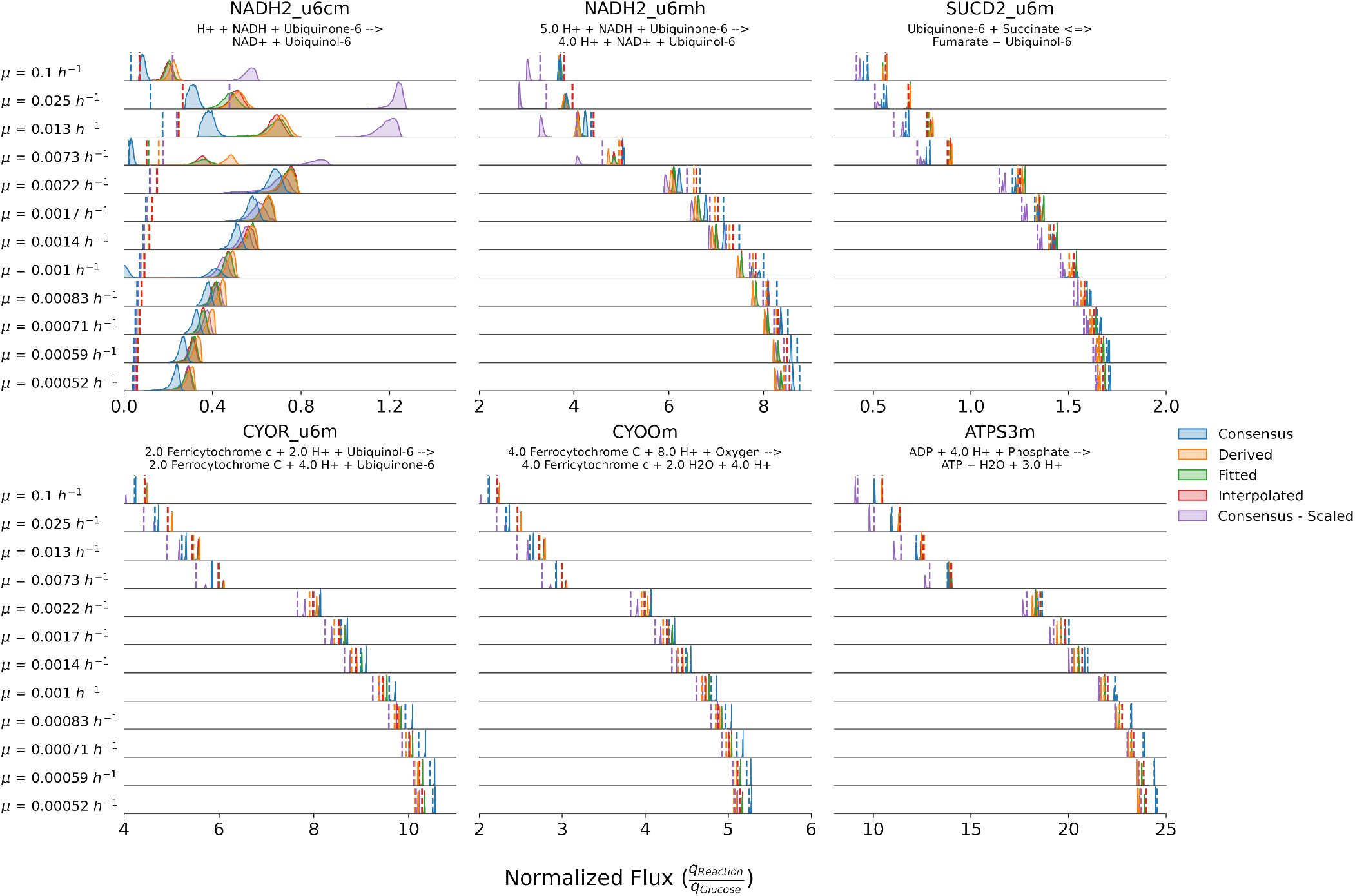
Glucose uptake normalised flux distributions for a number of reactions reactions involved in respiration and the electron transport chain. Dashed lines represent pFBA solutions, density plots are generated from flux sampling solutions. Different colours distinguish predictions generated using the different types of biomass equation, and their respective energetic parameters. NADH2_u6cm = external, alternative NADH dehydrogenase; NADH2_u6mh = complex I; SUCD2_u6m = complex II; CYOR_u6m = complex III; CYOOm = complex IV; ATPS3m = ATP synthase

**FIGURE 6.**
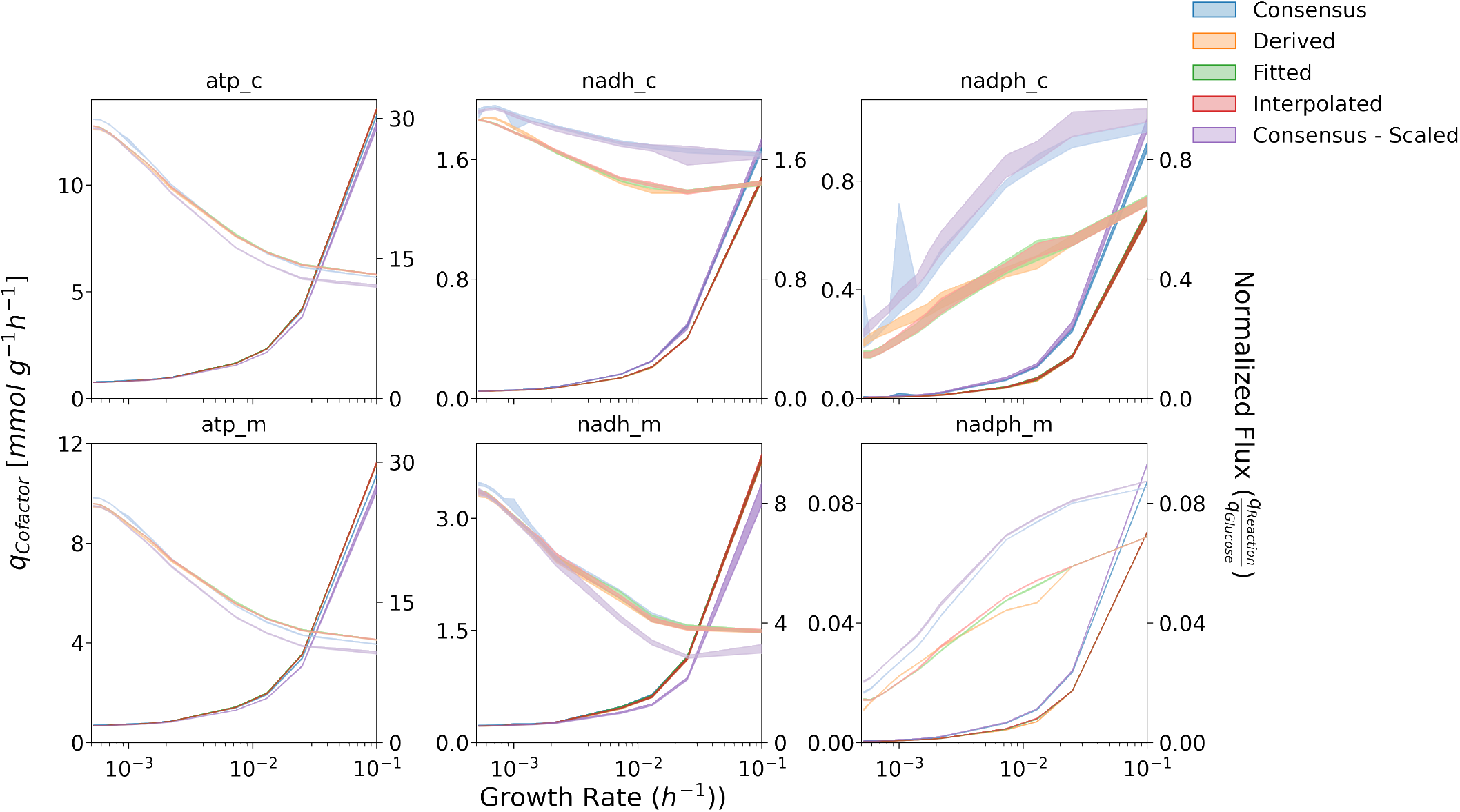
Total (bold) and glucose uptake normalised (faint) production rates of ATP (left), NADH (central) or NADPH (right) in the cytosol (top) or mitochondria (bottom) across different growth rates. Cytosolic ATP includes the ATP exported from the mitochondria. Shaded areas represent the 95% confidence interval from all sampled flux distributions at each sampling point.

Interestingly, both pFBA and sampling predicted a flux was carried by the external alternative NADH dehydrogenase (NADH2_u6cm) at 0.1 h^−1^, which increased between 0.025 h^−1^ and 0.0022 h^−1^, before decreasing at slower growth rates. No flux was observed for the internal dehydrogenase for either method. Both enzymes are present in *i*MT1026v3 based on homology to *S. cerevisiae* sequences, however, it is unclear to date which genes encode for which activity in *K. phaffii* (Tomàs-Gamisans et al. 2016; Valli et al. 2020).

The total, and glucose-relative, biomass-specific rates of production for cytosolic ATP, NADH and NADPH were very similar with the three methods used for biomass equation generation. The total biomass-specific rates of production for cytosolic NADH and NADPH decreased at slow growth rates (Figure 6). The flux yield for total NADH production increased slightly from ^∼^1.3 at 0.1 h^−1^ to 1.8 mmol_NADH_ mmol_glucose_^−1^ at 0.00059 h^−1^. In contrast, NADPH production decreased from ∼0.6 at 0.1 h^−1^ to 0.25 mmol_NADPH_ mmol_glucose_^−1^ at 0.00059 h^−1^. There were small changes in the relative contribution of each production source to total cofactor production (Table S7). NADPH supply remained relatively stable and was almost solely derived from the PPP, only a small amount was coming from folate metabolism. For NADH, there was increase in the contribution of glycolysis to total production when cells approached slower growth, accompanied with a decreased contribution from the TCA cycle. On the other hand, TCA cycle and oxidative phosphorylation contributed more to total ATP production at slower growth rates. At 0.1 h^−1^, ∼18 % of produced ATP was from glycolysis, whilst at slow growth rates this was reduced to ∼12 %. Thus, the vast majority of changes within metabolism are aligned with the increasing proportion of glucose committed to the TCA cycle and ATP generation.

### Biomass equation dependent flux predictions

Whilst the general pattern of changes were shared between the different biomass equations used, there was a dependency of flux predictions on the method used for biomass equation generation. We performed pairwise comparisons using the sampled flux distributions at the same growth rate but generated using different biomass equations. On average, 71 % of reactions were significantly differently distributed (MannWhitney U, MWU) and 44 % of all reactions demonstrated a large effect size (stochastic dominance of one distribution over the other) (Vargha, Delaney, & Vargha 2000). Although, it is worth noting that despite the fitted and interpolated methods using the same biomass equation and energetic parameters at 0.1 h^−1^ and 0.025 h^−1^, random sampling and the large sample size leads to 44 % and 49 % of reactions being determined as significantly differently distributed (MWU) and ∼6 % and 12 % had a large effect size, respectively.

The fitted and interpolated equations produced more similar fluxes across all growth rates with an average of 32 % reactions demonstrating a large effect. This was expected based on the energetic parameters NGAM and GAM being the same for the two methods, and the stoichiometry being very similar (Figure S2). The results generated with the derived equations differed in comparison to the fitted and interpolated equations across all growth rates, with ∼43 % of reactions demonstrating a large effect size. Compared to the fitted and interpolated equations, the stoichiometry of the derived equations differed more at slower growth rates (Figure S2). Additionally, at slower growth rates the slightly lower NGAM of the derived equations has a greater effect. The consensus and scaled consensus equations differed both to each other and to all three methods, with an average of over 45 % of reactions differing in each comparison with a large effect size. The largest differences occurred at higher growth rates, where by definition of the Pirt equation the stoichiometry of the biomass equation exerts the greatest effect on substrate usage and therefore metabolic fluxes.

We performed enrichment analysis on the pathways containing more than 5 reactions that had a large effect size. In over 70 % of comparisons, there was significant enrichment of pathways including: sterol metabolism, histidine metabolism, sphingolipid metabolism, fatty acid biosynthesis and the pentose phosphate pathway. The TCA cycle, glycolysis and oxidative phosphorylation were significantly enriched in 69 %,58 % and 1 % of comparisons, respectively.

Despite being determined at the same growth rate of 0.1 h^−1^ (Carnicer et al. 2009; Tomàs-Gamisans et al. 2016), absolute and relative differences were observed between the consensus equation, the scaled consensus our 0.1 h^−1^ specific equations, particularly around the oxidative PPP and glycolysis entry (Table S6). The median flux ratio for the consensus and scaled consensus equation entering glycolysis through PGI was ∼38 % and ∼36 %, and entering PPP via G6PDH2 was ∼44 % and ∼45 %, respectively. Our 0.1 h^−1^ specific equations suggested ∼44 % entered glycolysis and ∼31 % entered PPP (Figure 3). This difference could be explained by the reduced carbohydrate content of the consensus equation, as the majority of metabolites contributing to the carbohydrate macromolecule are derived from glucose-6-phosphate (Figure S3). There were also differences in other metabolic pathways. Between 0.1 h−1 to 0.0073 h^−1^, the flux yield predictions with the growth rate specific equations in comparison to the consensus equations were: lower for lower glycolysis (phosphofructokinase, pyruvate kinase, pyruvate carboxylase), oxidative phosphorylation and the malate-aspartate shuttle (complex I); but higher for a number of reactions in the TCA cycle as well as the external alternative NADH dehydrogenase (Figure 3 and Figure 5). At lower growth rates, the growth rate specific equations and consensus equations predicted very similar results, as the dominant constraints are substrate availability and the NGAM (Table S6).

The differences in predicted fluxes extended into the total and glucose normalised cofactor production rates (Figure 6), although a very similar pattern of changes was observed and the production sources were very similar (Table S7). At higher growth rates, the total and glucose normalised production rates of cytosolic and mitochondrial NADH and NADPH were higher than with the other equations, whereas ATP was much the same.

## DISCUSSION

### Maintenance requirements of *K. phaffii*

Here, we used CBM methods to quantitatively determine the metabolic rewiring of a recombinant protein-producing *K. phaffii* strain during the transition to glucose-limited nearzero growth rates. The *m*_s_ of *K. phaffii* is growth-rate dependent, dropping 3-fold between 0.06 h^−1^ and 0.03 h^−1^ (Rebnegger et al. 2023 2016). Here, we observed that between 0.00052 h^−1^ to 0.1 h^−1^, growth-rates could be reliably predicted with a constant NGAM and GAM. Small differences in these parameters existed between different biomass equations, but with the equations generated in this study, an NGAM of 0.65 mmol_ATP_ g_CDW_^−1^ h^−1^ and a GAM of 108 mmol_ATP_ g_CDW_^−1^ were determined. By definition of the Pirt equation, as *q*_s_ decreases, NGAM becomes dominant which on a metabolic level leads to a rewiring pattern characterised by an increase in the yield of ATP from glucose.

The consensus biomass equation of *i*MT1026v3 has been utilised for metabolic flux predictions by both Torres, Saa, Albiol, Ferrer, and Agosin (2019) and Tomàs-Gamisans et al. (2016). For the consensus biomass equation, the GAM was originally determined as 72 mmol_ATP_ g_CDW_^−1^ and the NGAM as 2.5 mmol_ATP_ g_CDW_^−1^ h^−1^ in (Tomàs-Gamisans et al. 2016). Our estimates for GAM are higher, whilst our NGAM estimates are lower. However, with the consensus equation at a growth-rate of 0.1 h^−1^, our *q*_ATP_ of 10.2 mmol_ATP_ g_CDW_^−1^ is similar to the 9.7 mmol_ATP_ g_CDW_^−1^ when using the original energetic parameters of *i*MT1026v3 (Tomàs-Gamisans et al. 2016). Torres et al. (2019) incorporated the earlier *m*_s_ estimate of a non-producing strain of *K. phaffii* from glucose-limited retentostats to predict an NGAM of 0.55 mmol_ATP_ g_CDW_^−1^ h^−1^, which is slightly lower than our estimate. Additionally, our GAM estimates are at the upper end of the range of GAM determined at different oxygenation levels, which ranged from 66 mmol_ATP_ g_CDW_^−1^ to 95 mmol_ATP_ g_CDW_^−1^ (Torres et al. 2019). Nevertheless, the NGAM of *K. phaffii* is ∼60 % lower than reported for *S. cerevisiae* in similar conditions (Dinh & Maranas 2022).

However, it is important to highlight that below a growth rate of 0.0022 h^−1^ we assumed a 50 % relative error to the carbon dioxide production rates, and across all growth rates a relative error of 50 % for oxygen uptake rates. We assumed these errors based on large variation in oxygen transfer rates between the replicate retentostat cultivations, and a lack of sensitivity of the gas sampler at very low glucose feed rates. Changes in accurate gas exchanges likely affect flux predictions, particularly involving energy metabolism; thus, their influence at near-zero growth rates needs to be further investigated. Yet, across the almost 200-fold range of growth-rates investigated here, we found that a static NGAM and GAM could be used to accurately predict growth-rates below 0.1 h^−1^.

### Coordinated metabolic changes increase the efficiency of ATP generation

We observed a negative correlation between the growthrate/glucose uptake rate and the yield of ATP per glucose molecule. This increase was mediated by changes in central carbon metabolism and in the usage of components of the respiratory chain. At slower growth rates, a smaller proportion of glucose derived flux entered the PPP, and was instead directed through glycolysis towards the TCA cycle. This reduction in PPP entry reduces the NADPH yield, which is considered important for protein synthesis and biomass generation (Heyland et al. 2010). Interestingly, as the PPP flux gets lower, the portion of NADPH production attributable to folate metaoblism, specifically methylene-THF dehydrogenase. increased. The metabolic rewiring was partly reflected in the transcriptome adaptation to near-zero growth (Rebnegger et al. 2023). Genes encoding for the three subunits of PFK (*PFK1, PFK2, PFK300*), citrate synthase *CIT1*, and aconitase *ACO1* were upregulated at near-zero growth. Yet, a number of genes in the pentose phosphate pathway, gluconeogensis and alternative carbon metabolism were also upregulated.

The increasing flux yield of all components of the respiratory chain further highlights the increasing efficiency of ATP production from glucose at slower growth rates. Under energy limitation, this increase in efficiency seems logical, whilst the activity of a non-proton translocating, and energy inefficient alternative NADH dehydrogenase seems counterintuitive. Juergens et al. (2020) hypothesised that an alternative NADH dehydrogenase was active in the related methylotrophic yeast *O. polymorpha*, based on the complex I-independent growth rate dependency of *m*_s_ (Juergens et al. 2020). Juergens et al. (2021) later identified an internal alternative dehydrogenase, and additionally observed that both an external alternative dehydrogenase and shuttle mechanism for the respiration of cytosolic NADH must exist. Our findings suggest that in *K. phaffii* an external alternative NADH dehydrogenase is active at slow growth rates, and that the aspartate-malate shuttle allows for the respiration of cytosolic NADH via complex I. Through protein sequence homology to *S. cerevisiae*, two alternative dehydrogenases (PP7435_Chr1–1084 and PP7435_Chr3-0399) are annotated in *K. phaffii*, in addition to an alternative oxidase (AOX100, PP7435_Chr3-0805) (Kern et al. 2007; Mar González-Barroso et al. 2006; Valli et al. 2020). Based on this, (Tomàs-Gamisans et al. 2016) defined both an internal and external alternative dehydrogenase in *i*MT1026v3, both of which were unconstrained, and hence able to carry flux in our analysis.

It is yet to be determined whether *K. phaffii* has both an external and internal NADH dehydrogenase, although Mar González-Barroso et al. (2006) demonstrated that isolated mitochondria from *K. phaffii* can oxidise external NADH, suggesting the presence of an external NADH dehydrogenase. At slower growth rates, the transcript levels of PP7435_Chr1–1084 are upregulated, whilst at faster growthrates the transcript levels of PP7435_Chr3-0399 and the alternative oxidase are upregulated (Kern et al. 2007; Rebnegger et al. 2014 2023 2016). Proteomics data of slow growing *K. phaffii* is not available, but in rich glucose conditions only PP7435_Chr1–1084 was quantified (Valli et al. 2020). Based on this correlation, Valli et al. (2020) suggested that PP7435_Chr1-1084 was an internal dehydrogenase (NDI1) and that PP7435_Chr3-0399 an external (NDE2). In comparison to the chemostat at 0.1 h^−1^, PP7435_Chr1-1084 was upregulated at all sampling points during the retentostat cultivation, but reduced at the slowest growth rate; following a similar trajectory to the predicted flux yield (Figure 5) (Rebnegger et al. 2023). Experimental work is required to verify our hypothesis; however, based on the transcript levels and flux distributions we suggest that PP7435_Chr1–1084 is a unique external NADH dehydrogenase, and that PP7435_Chr3-0399 is likely an internal NADH dehydrogenase.

Despite being energetically less efficient, the increase in alternative dehydrogenase utilisation with increasing substrate likely reflects the lower production costs of a single subunit protein over the 41 subunit complex I of *K. phaffii* (Bridges, Fearnley, & Hirst 2010). In conditions of glucose excess, Crabtree-negative yeast maximise ATP yield by switching to energetically less efficient pathways, with lower protein production costs (Malina, Yu, Björkeroth, Kerkhoven, & Nielsen 2021). Furthermore, protein production costs have been implicated in the trade-off between growth rate and yield (Wortel, Noor, Ferris, Bruggeman, & Liebermeister 2018), and in overflow metabolism (Chen & Nielsen 2019).

To apply CBM methods, we assumed a steady-state through the retentostat cultivation which limits our analysis. The intracellular concentrations of many metabolites are dynamic and have been implicated in sensing metabolic flux (Hackett et al. 2016; Litsios, Ortega, Wit, & Heinemann 2018; Xu et al. 2012). Accounting for the dynamics of metabolite pools would enhance flux predictions, particularly at slow growth. However, absolute quantification of the intracellular metabolome is both difficult and expensive, especially in the context of retentostat cultivations. Accounting for the dynamic changes in the metabolome and proteome would enhance flux prediction and enable a deeper understanding of the metabolic rewiring at near-zero growth rates, as it has for probing the physiological changes in *E. coli* and *S. cerevisiae* (Dinh & Maranas 2022; Grigaitis et al. 2021; Lu et al. 2019; Oftadeh et al. 2021).

#### The value of context-dependent biomass equations

While all biomass equations predicted a similar pattern of metabolic rewiring, predicted flux distributions were sensitive to the biomass equation. This supports previous findings that an accurate and context-dependent biomass equation is important for predictive accuracy of a GSMM with CBM methods (Dikicioglu et al. 2015). Clear differences, particularly at higher growth rates, were observed around the initial branch points of central-carbon metabolism. At 0.1 h^−1^, the split flux ratio of glucose entering the PPP ranged between 43 % to 44 % and 44 % to 47 % when using the consensus and scaled consensus equation, respectively. Whereas for all three methods, the flux yield ranged 31 % to 33 %. Values between 25 % to 50 % have been reported a producing K. phaffii strain and can vary for a number of reasons, including: cultivation differences, strain background differences, recombinant-protein specific effects and strain engineering effects (Heyland et al. 2010; Jordà et al. 2012; Saitua, Torres, Pérez-Correa, & Agosin 2017; Torres et al. 2019). For example, splits up to 60 % have been reported for engineered *K. phaffii* strains (Nocon et al. 2016). The differences in predictions highlight the importance of an accurate biomass equation in determining metabolic fluxes with CBM methods, supporting previous findings (Dikicioglu et al. 2015; Dinh et al. 2022).

The consensus glucose biomass equation was generated using the average biomass composition from normoxic (21 % oxygen) 0.1 h^−1^ chemostats of both a X-33 control strain and a FAB antibody-fragment producing derivative cultivated at 25 °C (Carnicer et al. 2009; Tomàs-Gamisans et al. 2016). The strain used by Rebnegger et al. (2023) was a vHH-producing, flo8-deficient strain derived from the CBS2612 background and was cultivated at 30 °C. Significant physiological differences have been reported between the closely related strains of *K. phaffii*, including at the transcriptional level and in relation to their protein producing capacity (Brady et al. 2020; Offei et al. 2022). Our results demonstrate that a context-specific biomass equation can help identifying these condition specific effects; however, the determination of the biomass composition remains a time and cost intensive process.

Three approaches were described to represent the changing biomass composition during the retentostat cultivation. The equations generated with the interpolated method are similar to the approach used by Y. Liu et al. (2019), who used the biomass composition measured at the end of the retentostat cultivation to predict metabolic flux distributions of nitrogen- and phosphorous-limited *S. cerevisiae*. The equations generated with the derived method, originally used by Goffin et al. (2010), assume that newly generated biomass is the sole contributor to the changing biomass composition. The fitted, and subsequently the derived method, required the fitting of a number of functions to the biomass component concentration. As the stoichiometry of the fitted and interpolated methods were very similar, we concluded that the chosen functions provided a reasonable approximation of accumulation of the individual biomass components.

With more data points for the biomass glycogen content, we observed a reduction in the total glycogen concentration in the reactor, indicating consumption. The derived method was the only method that accounted for a consumption of glycogen between 0.0014 h^−1^ and 0.00052 h^−1^. In glucose-limited conditions, storage carbohydrates have been reported to accumulate, and upon starvation (which is inherently different to that studied here) are rapidly depleted (Weber et al. 2020). A reduction in the glycogen content has previously been observed at extremely slow growth rates in *S. cerevisiae* (L. G. Boender et al. 2011; Vos et al. 2016). However, despite comparing accuracy of growth rate predictions (which were extremely similar) and the ability of the different methods to account for circumstances such as glycogen consumption, it is difficult to determine which biomass equation generation method is optimal without comparison to verified metabolic fluxes e.g. from labelling experiments.

## CONCLUSION

Through quantitatively analysing the transition to near-zero growth rates of *K. phaffii*, we could enhance our understanding of the metabolic adaptations to severe energy source limitation. We extended a regression model describing glucose-limited retentostat cultivations and generated a number of growth rate specific biomass equations of a protein-producing *K. phaffii* strain. Constraint-based modelling methods revealed a pattern of metabolic rewiring in the central carbon metabolism and respiratory chain of *K. phaffii*, largely attributable to the increasing efficiency of ATP generation at slower growth rates. A similar trend and estimated energetic parameters were predicted with both a consensus glucose biomass equation, and those generated in this study. Differences in flux predictions between the equations did exist and likely reflect strain and condition specific differences, which will be important for guiding interventions in strain engineering and bioprocess design to uncouple protein production from growth in *K. phaffii*.

## METHODS

Executable notebooks and necessary data for reproduction are available on GitHub at www.github.com/bcoltman/Kphaffii_NearZero. All analysis was performed in Python v3.7.11. Key libraries used include: COBRApy v0.24.0 (Ebrahim, Lerman, Palsson, & Hyduke 2013) for all CBM methods and model adaptations, Pandas v1.3.4 (Reback et al. 2021), NumPy v1.21.2 (Harris et al. 2020), SciPy v1.7.3 (Virtanen et al. 2020) and Scikit-learn v1.0.2(Pedregosa et al. 2011).

### Retentostat regression model

The series of ODEs describing the retentostat cultivation (Rebnegger et al. 2016) were translated into Python and adapted according to the results section. An optimal *m*_s_ was separately determined for each cultivation through minimising the weighted sum-of-squares between regression model predictions and experimental data. The integrated concentrations and derived rates were used at later points in this analysis.

### GSMM updates and biomass equation generation

The base GSMM used throughout this study was *i*MT1026v3, downloaded from https://www.ebi.ac.uk/biomodels/ MODEL1612130000. Slight modifications were made, including the incorporation of reactions for recombinant protein production (Supplementary Methods S1.1). All data on biomass compositions and cultivation parameters were extracted from Rebnegger et al. (2023). Data was available at multiple sampling points and included: carbohydrate composition, amino acid composition and lipid composition, amongst others. Additional static compositions for some components not measured in this study were assumed, as in previous *K. phaffii* biomass compositions (Tomàs-Gamisans et al. 2016) (Supplementary Methods S1.2). Details for how the biomass equation generation methods differ can be found in the results section (Generation of growth rate dependent biomass equations).

The sub-compositions of the macromolecules were converted into mmol g_CDW_^−1^, and the macromolecule classes represented as g g_CDW_^−1^. We did not include the energetic costs for polymerisation of the macromolecules, choosing to include this in GAM calculations as they represent only a small fraction of the GAM. The new model, *i*MT1026-NZ, containing the biomass equations and updates is available at www.github.com/bcoltman/Kphaffii_NearZero and https://www.ebi.ac.uk/biomodels/TO_UPDATE.

### GSMM constraints

Biomass-specific glucose uptake rates *q*_*s*_, recombinant protein secretion rates *q*_*p*_ and growth rates *μ* for each individual cultivation were derived from the regression model. Oxygen uptake rate (OUR) and carbon production rate (CPR) were calculated from mass balancing, before a Gaussian process regressor with a WhiteNoise kernel was fitted to the data, enabling propagation of error to the calculated biomass-specific exchange rates 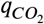 and 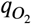 S1.3. Due to large variance observed in the OUR, at each sampling point the upper and lower bound were set to ± 50 % of the average 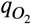 of the three cultivations. For 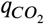, at sampling points above 0.0022 h^−1^, we set the bounds to ± 1 SD, determined from the Gaussian process regressor and propagated as above. Below 0.0022 h^−1^ we assumed a lack of sensitivity of the gas analyser and set the bounds to ± 50 % of the average of the cultivations. Commonly secreted metabolites including arabitol, ethanol and citrate were not detected in the reactor media, thus their secretion was blocked. Where appropriate, a sink reaction for glycogen consumption was added to the model, with an upper bound set to the timepoint dependent specific consumption rate. The consumption rate was calculated from the product of the growth rate and the stoichiometric coefficient from the derived biomass equation. Phosphate, ammonium and sulphate exchanges were not constrained as they were available in excess.

### Energetic parameter fitting

At each sampling point, we constrained the metabolic model with calculated exchange rates *q*_*s*_, 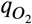, 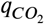 and *q*_*p*_, setting upper and lower bounds as stated above. Then, the biomass formation reaction was fixed to the respective regression model derived *μ* and using the ATPM reaction (NGAM) maximised via FBA to calculate *q*_*AT P*_ . A relative weighted linear fit of *q*_*AT P*_ vs *μ* was performed, and the intercept used as NGAM. To calculate GAM, we updated each biomass equation with a GAM in the range 0 mmol_ATP_ g_CDW_ to 200 mmol_ATP_ g_CDW_, then constrained the model and using FBA calculated the maximum growth rate at every sampling point. The optimal GAM was determined as the value that minimised the error-weighted sum of squares between the predicted growth rates and regression model derived growth rates.

### Comparative flux distribution analysis

To generate flux distributions for comparisons, we used pFBA and flux sampling. For pFBA, at each sampling point, the model was constrained with the respective cultivation parameters and the different biomass equations before the growth reaction reaction was maximised. For flux sampling, the upper bounds on the growth reaction were set to 105 % of the regression model predicted growth rate, while lower bounds set to 95 % of the lowest of either the maximal FBA growth rate or the regression model growth rate. Then, to generate the solutions, the following steps were performed:

i. When constrained, reactions carrying 0 flux at a fraction of optimality of 0 were identified through flux variability analysis (FVA), and subsequently removed from the model. No tolerance value was used due to model infeasiblity after the removal of some reactions.
ii. Flux sampling was performed using the OptGp sampler (Megchelenbrink et al. 2014) within COBRApy, 4 chains of 1,250,000 samples were generated, of which 1,250 were stored, representing a thinning rate of 1,000 that was determined as optimal for sampler convergence(see S2.1).
iii. Thermodynamically infeasible loops in sampled flux distributions were removed through the COBRApy implementation of the CycleFreeFlux algorithm (Desouki, Jarre, Gelius-Dietrich, & Lercher 2015), which converts a solution containing loops to the closest thermodynamically feasible solution.

The convergence of the sampler was determined as below, and the chains were merged together for further analysis and figure generation. Escher (King et al. 2015) was used for visualisation of metabolic pathways. The total formation rates of a number of co-factors were assessed for every sampled flux distribution by summing all of the positive products of the metabolite stoichiometric matrix and reaction flux vectors. To determine the sources of metabolite production, the subsystems defined within *i*MT1026v3 were used.

### Assessing sample convergence and optimal sample thinning

Ensuring convergence of the sampler is important for ensuring an accurate characterisation of the solution space. A converged sample chain should represent the properties of true solutions obtained from an infinite number of samples. Different diagnostic tests can be used to assess this convergence, both on a within-chain and between-chain basis. To assess convergence of our sampled chains, we implemented a number of diagnostic tests, based on sampling recommendations (Herrmann et al. 2019; Kumar, Carroll, Hartikainen, & Martin 2019; Vehtari, Gelman, Simpson, Carpenter, & Bürkner 2021). Three different general diagnostics were used:

i. The Geweke diagnostic was used to assess the ergodicity (stationarity) of a chain by comparing the first 10 % of a chain with 100 intervals of the final 50 % i.e determining of the means of the two subsets are different. A chain was deemed to have failed if the calculated *z* score for any of the investigated intervals was > 1.28.
ii. The rank normalized folded-split-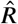 (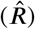) assesses a lack of convergence of a set of chains by taking into account between- and within-chain variance i.e. determining if the chains are mixed well. In line with (Vehtari et al. 2021) we deemed the sample converged if the calculated 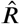 was less than 1.01.
iii. The Effective sample size (ESS) is a scale-free measure of information, representing the number of independent draws with the same estimation power as the dependent sample obtained by flux sampling i.e. determining the effects of autocorrelation within chains on the uncertainty of estimates. As recommended (Vehtari et al. 2021), the total bulk ESS across all chains was required to be above 400.

The flux sampling distribution of a reaction was deemed to have converged if the criteria outlined for 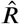 and ESS were met. The optimal thinning rate to achieve convergence of the sampled chains was assessed using the cultivation values for the chemostat at 0.1 h^−1^. The 0.1 h^−1^ biomass equation of the three methods, as well as the consensus and scaled consensus biomass equations were tested using the cultivation data at 0.1 h^−1^. 4 chains were generated and a total of 5,000, 50,000, 500,000 and 5,000,000 samples were generated, 5,000 of which were stored, representing thinning factors of 1, 10, 100 and 1,000, respectively.

## AUTHOR CONTRIBUTION

Conceptualisation: B.L.C., C.R., B.G., J.Z.

Methodology and Formal Analysis: B.L.C. and C.R.

Data Acquisition and Curation: B.L.C. and C.R.

Software Implementation: B.L.C.

Visualisation: B.L.C.

Writing – Original Draft: B.L.C. and J.Z.

Writing – Review & Editing: all

Funding Acquisition: B.G.

## DATA AVAILABILITY

*i*MT1026v3 from (Tomàs-Gamisans et al. 2018) is publicly available on BioModels (Malik-Sheriff et al. 2020) with ascension ID MODEL1612130000 https://www.ebi.ac.uk/biomodels/MODEL1612130000.

The model generated in this analysis is available at www.github.com/bcoltman/Kphaffii_NearZero and https://www.ebi.ac.uk/biomodels/TO_UPDATE.

## CODE AVAILABILITY

Jupyter Notebooks and source files of this analysis are available at www.github.com/bcoltman/Kphaffii_NearZero.

## ACKNOWLEDGEMENTS

This work has been funded by the Austrian Federal Ministry of Digital and Economic Affairs (BMDW), the Nationalstiftung für Forschung, Technologie und Entwicklung, and Lonza AG through the Christian Doppler Research Association. Further support was given by the Federal Ministry of Digital and Economic Affairs (BMDW), the Federal Ministry of Traffic, Innovation and Technology (bmvit), the Styrian Business Promotion Agency SFG, the Standortagentur Tirol, the Government of Lower Austria, the Business Agency Vienna and BOKU through the COMET Funding Program managed by the Austrian Research Promotion Agency FFG within the COMET center: acib – Next Generation Bioproduction.

## References

Binai, N. A., Bisschops, M. M. M., van Breukelen, B., Mohammed, S., Loeff, L., Pronk, J. T., … Slijper, M. (2014). Proteome Adaptation of Saccharomyces cerevisiae to Severe Calorie Restriction in Retentostat Cultures. Journal of Proteome Research, 13(8), 3542–3553. doi: 10.1021/pr5003388

Bisschops, M. M. M., Luttik, M. A. H., Doerr, A., Verheijen, P. J. T., Bruggeman, F., Pronk, J. T., & Daran-Lapujade, P. (2017). Extreme calorie restriction in yeast retentostats induces uniform non-quiescent growth arrest. Biochimica et Biophysica Acta (BBA) - Molecular Cell Research, 1864(1), 231–242. doi: 10.1016/j.bbamcr.2016.11.002

Boender, L. G., van Maris, A. J., de Hulster, E. A., Almering, M. J., van der Klei, I. J., Veenhuis, M., … Daran-Lapujade, P. (2011). Cellular responses of Saccharomyces cerevisiae at near-zero growth rates: Transcriptome analysis of anaerobic retentostat cultures. FEMS Yeast Research, 11(8), 603–620. doi: 10.1111/j.1567-1364.2011.00750.x

Boender, L. G. M., de Hulster, E. A. F., van Maris, A. J. A., Daran-Lapujade, P. A. S., & Pronk, J. T. (2009). Quantitative Physiology of Saccharomyces cerevisiae at Near-Zero Specific Growth Rates. Applied and Environmental Microbiology, 75(17), 5607–5614. doi: 10.1128/AEM.00429-09

Brady, J. R., Whittaker, C. A., Tan, M. C., Kristensen II, D. L., Ma, D., Dalvie, N. C., … Love, J. C. (2020). Comparative genome-scale analysis of Pichia pastoris variants informs selection of an optimal base strain. Biotechnology and Bioengineering, 117(2), 543–555. doi: 10.1002/bit.27209

Bridges, H. R., Fearnley, I. M., & Hirst, J. (2010). The Subunit Composition of Mitochondrial NADH:Ubiquinone Oxidoreductase (Complex I) From Pichia pastoris. Molecular & Cellular Proteomics, 9(10), 2318–2326. doi: 10.1074/mcp.M110.001255

Buchetics, M., Dragosits, M., Maurer, M., Rebnegger, C., Porro, D., Sauer, M., … Mattanovich, D. (2011). Reverse engineering of protein secretion by uncoupling of cell cycle phases from growth. Biotechnology and Bioengineering, 108(10), 2403–2412. doi: 10.1002/bit.23198

Carnicer, M., Baumann, K., Töplitz, I., Sánchez-Ferrando, F., Mattanovich, D., Ferrer, P., & Albiol, J. (2009). Macromolecular and elemental composition analysis and extracellular metabolite balances of Pichia pastoris growing at different oxygen levels. Microbial Cell Factories, 8(1), 65. doi: 10.1186/1475-2859-8-65

Chan, S. H. J., Cai, J., Wang, L., Simons-Senftle, M. N., & Maranas, C. D. (2017). Standardizing biomass reactions and ensuring complete mass balance in genome-scale metabolic models. Bioinformatics, 33(22), 3603–3609. doi: 10.1093/bioinformatics/btx453

Chen, Y., & Nielsen, J. (2019). Energy metabolism controls phenotypes by protein efficiency and allocation. Proceedings of the National Academy of Sciences, 116(35), 17592–17597. doi: 10.1073/pnas.1906569116

Cregg, J. M., Cereghino, J. L., Shi, J., & Higgins, D. R. (2000). Recombinant protein expression in Pichia pastoris. Molecular Biotechnology, 16(1), 23–52. doi: 10.1385/MB:16:1:23

Desouki, A. A., Jarre, F., Gelius-Dietrich, G., & Lercher, M. J. (2015). CycleFreeFlux: Efficient removal of thermodynamically infeasible loops from flux distributions. Bioinformatics, 31(13), 2159–2165. doi: 10.1093/bioin-formatics/btv096

Dikicioglu, D., Kırdar, B., & Oliver, S. G. (2015). Biomass composition: The “elephant in the room” of metabolic modelling. Metabolomics, 11(6), 1690–1701. doi: 10.1007/s11306-015-0819-2

Dinh, H. V., & Maranas, C. D. (2022). Evaluating proteome allocation of Saccharomyces cerevisiae phenotypes with resource balance analysis. bioRxiv. doi: 10.1101/2022.09.20.508694

Dinh, H. V., Sarkar, D., & Maranas, C. D. (2022). Quantifying the propagation of parametric uncertainty on flux balance analysis. Metabolic Engineering, 69, 26–39. doi: 10.1016/j.ymben.2021.10.012

Ebrahim, A., Lerman, J. A., Palsson, B. O., & Hyduke, D. R. (2013). COBRApy: COnstraints-Based Reconstruction and Analysis for Python. BMC Systems Biology, 7(1), 74. doi: 10.1186/1752-0509-7-74

Garcia-Ortega, X., Adelantado, N., Ferrer, P., Montesinos, J. L., & Valero, F. (2016). A step forward to improve recombinant protein production in Pichia pastoris: From specific growth rate effect on protein secretion to carbon-starving conditions as advanced strategy. Process Biochemistry, 51(6), 681–691. doi: 10.1016/j.procbio.2016.02.018

Gasser, B., Prielhofer, R., Marx, H., Maurer, M., Nocon, J., Steiger, M., … Mattanovich, D. (2013). Pichia pastoris: Protein production host and model organism for biomedical research. Future Microbiology, 8(2), 191–208. doi: 10.2217/fmb.12.133

Goffin, P., van de Bunt, B., Giovane, M., Leveau, J. H. J., Höppener-Ogawa, S., Teusink, B., & Hugenholtz, J. (2010). Understanding the physiology of Lactobacillus plantarum at zero growth. Molecular Systems Biology, 6(1), 413. doi: 10.1038/msb.2010.67

Grigaitis, P., Olivier, B. G., Fiedler, T., Teusink, B., Kummer, U., & Veith, N. (2021). Protein cost allocation explains metabolic strategies in Escherichia coli. Journal of Biotechnology, 327, 54–63. doi: 10.1016/j.jbiotec.2020.11.003

Grillitsch, K., Tarazona, P., Klug, L., Wriessnegger, T., Zellnig, G., Leitner, E., … Daum, G. (2014). Isolation and characterization of the plasma membrane from the yeast Pichia pastoris. Biochimica et Biophysica Acta (BBA) - Biomembranes, 1838(7), 1889–1897. doi: 10.1016/j.bbamem.2014.03.012

Hackett, S. R., Zanotelli, V. R. T., Xu, W., Goya, J., Park, J. O., Perlman, D. H., … Rabinowitz, J. D. (2016). Systems-level analysis of mechanisms regulating yeast metabolic flux. Science, 354(6311), aaf2786–aaf2786. doi: 10.1126/science.aaf2786

Harris, C. R., Millman, K. J., van der Walt, S. J., Gommers, R., Virtanen, P., Cournapeau, D., … Oliphant, T. E. (2020). Array programming with NumPy. Nature, 585(7825), 357–362. doi: 10.1038/s41586-020-2649-2

Hastings, J., Owen, G., Dekker, A., Ennis, M., Kale, N., Muthukrishnan, V., … Steinbeck, C. (2016). ChEBI in 2016: Improved services and an expanding collection of metabolites. Nucleic acids research, 44(D1), D1214–9. doi: 10.1093/nar/gkv1031

Herrmann, H. A., Dyson, B. C., Vass, L., Johnson, G. N., & Schwartz, J.-M. (2019). Flux sampling is a powerful tool to study metabolism under changing environmental conditions. npj Systems Biology and Applications, 5(1), 1–8. doi: 10.1038/s41540-019-0109-0

Heyland, J., Fu, J., Blank, L. M., & Schmid, A. (2010). Quantitative physiology of Pichia pastoris during glucose-limited high-cell density fed-batch cultivation for recombinant protein production. Biotechnology and Bioengineering, 107(2), 357–368. doi: 10.1002/bit.22836

Jordà, J., Jouhten, P., Cámara, E., Maaheimo, H., Albiol, J., & Ferrer, P. (2012). Metabolic flux profiling of recombinant protein secreting Pichia pastoris growing on glucose:methanol mixtures. Microbial Cell Factories, 11(1), 57. doi: 10.1186/1475-2859-11-57

Juergens, H., Hakkaart, X. D. V., Bras, J. E., Vente, A., Wu, L., Benjamin, K. R., … Mans, R. (2020). Contribution of Complex I NADH Dehydrogenase to Respiratory Energy Coupling in Glucose-Grown Cultures of Ogataea parapolymorpha. Applied and Environmental Microbiology, 86(15), e00678–20. doi: 10.1128/AEM.00678-20

Juergens, H., Mielgo-Gómez, Á., Godoy-Hernández, A., ter Horst, J., Nijenhuis, J. M., McMillan, D. G. G., & Mans, R. (2021). Physiological relevance, localization and substrate specificity of the alternative (type II) mitochondrial NADH dehydrogenases of Ogataea parapolymorpha. bioRxiv. doi: 10.1101/2021.04.28.441406

Kern, A., Hartner, F. S., Freigassner, M., Spielhofer, J., Rumpf, C., Leitner, L., … Glieder, A. (2007). Pichia pastoris ‘just in time’ alternative respiration. Microbiology, 153(4), 1250–1260. doi: 10.1099/mic.0.2006/001404-0

King, Z. A., Dräger, A., Ebrahim, A., Sonnenschein, N., Lewis, N. E., & Palsson, B. O. (2015). Escher: A Web Application for Building, Sharing, and Embedding Data-Rich Visualizations of Biological Pathways. PLOS Computational Biology, 11(8), e1004321. doi: 10.1371/journal.pcbi.1004321

Kumar, R., Carroll, C., Hartikainen, A., & Martin, O. (2019). ArviZ a unified library for exploratory analysis of Bayesian models in Python. Journal of Open Source Software, 4(33), 1143. doi: 10.21105/joss.01143

Lewis, N. E., Hixson, K. K., Conrad, T. M., Lerman, J. A., Charusanti, P., Polpitiya, A. D., … Palsson, B. Ø. (2010). Omic data from evolved E. coli are consistent with computed optimal growth from genome-scale models. Molecular Systems Biology, 6(1), 390. doi: 10.1038/msb.2010.47

Litsios, A., Ortega, Á. D., Wit, E. C., & Heinemann, M. (2018). Metabolic-flux dependent regulation of microbial physiology. Current Opinion in Microbiology, 42, 71–78. doi: 10.1016/j.mib.2017.10.029

Liu, W.-C., Inwood, S., Gong, T., Sharma, A., Yu, L.-Y., & Zhu, P. (2019). Fed-batch high-cell-density fermentation strategies for Pichia pastoris growth and production. Critical Reviews in Biotechnology, 39(2), 258–271. doi: 10.1080/07388551.2018.1554620

Liu, Y., el Masoudi, A., Pronk, J. T., & van Gulik, W. M. (2019). Quantitative Physiology of Non-Energy-Limited Retentostat Cultures of Saccharomyces cerevisiae at Near-Zero Specific Growth Rates. Applied and Environmental Microbiology, 85(20), e01161–19. doi: 10.1128/AEM.01161-19

Liu, Y., Esen, O., Pronk, J. T., & van Gulik, W. M. (2021). Uncoupling growth and succinic acid production in an industrial Saccharomyces cerevisiae strain. Biotechnology and Bioengineering, 118(4), 1557–1567. doi: 10.1002/bit.27672

Liu, Z., Hou, J., Martínez, J. L., Petranovic, D., & Nielsen, J. (2013). Correlation of cell growth and heterologous protein production by Saccharomyces cerevisiae. Applied Microbiology and Biotechnology, 97(20), 8955–8962. doi: 10.1007/s00253-013-4715-2

Looser, V., Bruhlmann, B., Bumbak, F., Stenger, C., Costa, M., Camattari, A., … Kovar, K. (2015). Cultivation strategies to enhance productivity of Pichia pastoris: A review. Biotechnology Advances, 33(6, Part 2), 1177–1193. doi: 10.1016/j.biotechadv.2015.05.008

Lu, H., Li, F., Sánchez, B. J., Zhu, Z., Li, G., Domenzain, I., … Nielsen, J. (2019). A consensus S. cerevisiae metabolic model Yeast8 and its ecosystem for comprehensively probing cellular metabolism. Nature Communications, 10(1), 3586. doi: 10.1038

Malik-Sheriff, R. S., Glont, M., Nguyen, T. V. N., Tiwari, K., Roberts, M. G., Xavier, A., … Hermjakob, H. (2020). BioModels—15 years of sharing computational models in life science. Nucleic Acids Research, 48(D1), D407–D415. doi: 10.1093/nar/gkz1055

Malina, C., Yu, R., Björkeroth, J., Kerkhoven, E. J., & Nielsen, J. (2021). Adaptations in metabolism and protein translation give rise to the Crabtree effect in yeast. Proceedings of the National Academy of Sciences, 118(51), e2112836118. doi: 10.1073/pnas.2112836118

Mar González-Barroso, M., Ledesma, A., Lepper, S., Pérez-Magán, E., Zaragoza, P., & Rial, E. (2006). Isolation and bioenergetic characterization of mitochondria from Pichia pastoris. Yeast, 23(4), 307–313. doi: 10.1002/yea.1355

Mattanovich, D., Sauer, M., & Gasser, B. (2017). Industrial Microorganisms: Pichia pastoris. In Industrial Biotechnology (pp. 687–714). John Wiley & Sons, Ltd. doi: 10.1002/9783527807796.ch19

Maurer, M., Kühleitner, M., Gasser, B., & Mattanovich, D. (2006). Versatile modeling and optimization of fed batch processes for the production of secreted heterologous proteins with Pichia pastoris. Microbial Cell Factories, 5(1), 37. doi: 10.116/1475-2859-5-37

Megchelenbrink, W., Huynen, M., & Marchiori, E. (2014). optGpSampler: An Improved Tool for Uniformly Sampling the Solution-Space of Genome-Scale Metabolic Networks. PLOS ONE, 9(2), e86587. doi: 10.1371/journal.pone.0086587

Nocon, J., Steiger, M., Mairinger, T., Hohlweg, J., Rußmayer, H., Hann, S., … Mattanovich, D. (2016). Increasing pentose phosphate pathway flux enhances recombinant protein production in Pichia pastoris. Applied Microbiology and Biotechnology, 100(13), 5955–5963. doi: 10.1007/s00253-016-7363-5

Offei, B., Braun-Galleani, S., Venkatesh, A., Casey, W. T., O’Connor, K. E., Byrne, K. P., & Wolfe, K. H. (2022). Identification of genetic variants of the industrial yeast Komagataella phaffii (Pichia pastoris) that contribute to increased yields of secreted heterologous proteins. PLOS Biology, 20(12), e3001877. doi: 10.1371/journal.pbio.3001877

Oftadeh, O., Salvy, P., Masid, M., Curvat, M., Miskovic, L., & Hatzimanikatis, V. (2021). A genome-scale metabolic model of Saccharomyces cerevisiae that integrates expression constraints and reaction thermodynamics. Nature Communications, 12(1), 4790. doi: 10.1038/s41467-021-25158-6

Pedregosa, F., Varoquaux, G., Gramfort, A., Michel, V., Thirion, B., Grisel, O., … Duchesnay, É. (2011). Scikit-learn: Machine Learning in Python. Journal of Machine Learning Research, 12(85), 2825–2830.

Pirt, S. J. (1982). Maintenance energy: A general model for energy-limited and energy-sufficient growth. Archives of Microbiology, 133(4), 300–302. doi: 10.1007/BF00521294

Puxbaum, V., Gasser, B., & Mattanovich, D. (2016). The bud tip is the cellular hot spot of protein secretion in yeasts. Applied Microbiology and Biotechnology, 100(18), 8159–8168. doi: 10.1007/s00253-016-7674-6

Reback, J., jbrockmendel McKinney, W., den Bossche, J. V., Augspurger, T., Cloud, P., … Seabold, S. (2021). Pandas-dev/pandas: Pandas 1.3.4. Zenodo. doi: 10.5281/zenodo.5574486

Rebnegger, C., Graf, A. B., Valli, M., Steiger, M. G., Gasser, B., Maurer, M., & Mattanovich, D. (2014). In Pichia pastoris, growth rate regulates protein synthesis and secretion, mating and stress response. Biotechnology Journal, 9(4), 511–525. doi: 10.1002/biot.201300334

Rebnegger, C., Hann, S., Coltman, B., Mattanovich, D., Gasser, B., Kowarz, V., … Koellensperger, G. (2023). Protein production dynamics and physiological adaptation of recombinant Pichia pastoris at near-zero growth rates. bioRxiv.

Rebnegger, C., Vos, T., Graf, A. B., Valli, M., Pronk, J. T., Daran-Lapujade, P., & Mattanovich, D. (2016). Pichia pastoris Exhibits High Viability and a Low Maintenance Energy Requirement at Near-Zero Specific Growth Rates. Applied and Environmental Microbiology, 82(15), 4570–4583. doi: 10.1128/AEM.00638-16

Saitua, F., Torres, P., Pérez-Correa, J. R., & Agosin, E. (2017). Dynamic genome-scale metabolic modeling of the yeast Pichia pastoris. BMC Systems Biology, 11(1), 27. doi: 10.1186/s12918-017-0408-2

Schoeny, H., Rampler, E., Abiead, Y. E., Hildebrand, F., Zach, O., Hermann, G., & Koellensperger, G. (2021). A combined flow injection/reversed-phase chromatography–high-resolution mass spectrometry workflow for accurate absolute lipid quantification with 13C internal standards. Analyst, 146(8), 2591–2599. doi: 10.1039/D0AN02443K

Stephanopoulos, G., Aristidou, A. A., & Nielsen, J. (1998). Metabolic Engineering: Principles and Methodologies. Elsevier.

Széliová, D., Ruckerbauer, D. E., Galleguillos, S. N., Petersen, L. B., Natter, K., Hanscho, M., … Zanghellini, J. (2020). What CHO is made of: Variations in the biomass composition of Chinese hamster ovary cell lines. Metabolic Engineering, 61, 288–300. doi: 10.1016/j.ymben.2020.06.002

Széliová, D., Štor, J., Thiel, I., Weinguny, M., Hanscho, M., Lhota, G., … Rocha, I. (2021). Inclusion of maintenance energy improves the intracellular flux predictions of CHO. PLOS Computational Biology, 17(6), e1009022. doi: 10.1371/journal.pcbi.1009022

Thiele, I., & Palsson, B. Ø. (2010). A protocol for generating a high-quality genome-scale metabolic reconstruction. Nature protocols, 5(1), 93–121. doi: 10.1038/nprot.2009.203

Tomàs-Gamisans, M., Ferrer, P., & Albiol, J. (2016). Integration and Validation of the Genome-Scale Metabolic Models of Pichia pastoris: A Comprehensive Update of Protein Glycosylation Pathways, Lipid and Energy Metabolism. PLOS ONE, 11(1), e0148031. doi: 10.1371/journal.pone.0148031

Tomàs-Gamisans, M., Ferrer, P., & Albiol, J. (2018). Fine-tuning the P. pastoris iMT1026 genome-scale metabolic model for improved prediction of growth on methanol or glycerol as sole carbon sources. Microbial Biotechnology, 11(1), 224–237. doi: 10.1111/1751-7915.12871

Torres, P., Saa, P. A., Albiol, J., Ferrer, P., & Agosin, E. (2019). Contextualized genome-scale model unveils high-order metabolic effects of the specific growth rate and oxygenation level in recombinant Pichia pastoris. Metabolic Engineering Communications, e00103. doi: 10.1016/j.mec.2019.e00103

Valli, M., Grillitsch, K., Grünwald-Gruber, C., Tatto, N. E., Hrobath, B., Klug, L., … Mattanovich, D. (2020). A subcellular proteome atlas of the yeast Komagataella phaffii. FEMS Yeast Research, 20(1), foaa001. doi: 10.1093/femsyr/foaa001

Vargha, A., Delaney, H. D., & Vargha, A. (2000). A Critique and Improvement of the “CL” Common Language Effect Size Statistics of McGraw and Wong. Journal of Educational and Behavioral Statistics, 25(2), 101. doi: 10.2307/1165329

Vehtari, A., Gelman, A., Simpson, D., Carpenter, B., & Bürkner, P.-C. (2021). Rank-Normalization, Folding, and Localization: An Improved R^ for Assessing Convergence of MCMC (with Discussion). Bayesian Analysis, 16(2), 667–718. doi: 10.1214/20-BA1221

Virtanen, P., Gommers, R., Oliphant, T. E., Haberland, M., Reddy, T., Cournapeau, D., … Vázquez-Baeza, Y. (2020). SciPy 1.0: Fundamental algorithms for scientific computing in Python. Nature Methods, 17(3), 261–272. doi: 10.1038/s41592-019-0686-2

Vos, T., Hakkaart, X. D. V., de Hulster, E. A. F., van Maris, A. J. A., Pronk, J. T., & Daran-Lapujade, P. (2016). Maintenance-energy requirements and robustness of Saccharomyces cerevisiae at aerobic near-zero specific growth rates. Microbial Cell Factories, 15(1), 111. doi: 10.1186/s12934-016-0501-z

Wang, J., Wang, X., Shi, L., Qi, F., Zhang, P., Zhang, Y., … Cai, M. (2017). Methanol-Independent Protein Expression by AOX1 Promoter with trans-Acting Elements Engineering and Glucose-Glycerol-Shift Induction in Pichia pastoris. Scientific Reports, 7(1), 41850. doi: 10.1038/srep41850

Wanka, F., Arentshorst, M., Cairns, T. C., Jørgensen, T., Ram, A. F. J., & Meyer, V. (2016). Highly active promoters and native secretion signals for protein production during extremely low growth rates in Aspergillus niger. Microbial Cell Factories, 15(1), 1–12. doi: 10.1186/s12934-016-0543-2

Weber, C. A., Sekar, K., Tang, J. H., Warmer, P., Sauer, U., & Weis, K. (2020). β-Oxidation and autophagy are critical energy providers during acute glucose depletion in Saccharomyces cerevisiae. Proceedings of the National Academy of Sciences, 117(22), 12239–12248. doi: 10.1073/pnas.1913370117

Wortel, M. T., Noor, E., Ferris, M., Bruggeman, F. J., & Liebermeister, W. (2018). Metabolic enzyme cost explains variable trade-offs between microbial growth rate and yield. PLOS Computational Biology, 14(2), e1006010. doi: 10.1371/journal.pcbi.1006010

Xu, Y.-F., Zhao, X., Glass, D. S., Absalan, F., Perlman, D. H., Broach, J. R., & Rabinowitz, J. D. (2012). Regulation of yeast pyruvate kinase by ultrasensitive allostery independent of phosphorylation. Molecular cell, 48(1), 52–62. doi: 10.1016/j.molcel.2012.07.013

Yang, Z., & Zhang, Z. (2018). Engineering strategies for enhanced production of protein and bio-products in Pichia pastoris: A review. Biotechnology Advances, 36(1), 182–195. doi: 10.1016/j.biotechadv.2017.11.002

