## Supplementary Information for "Characterising the metabolic rewiring of extremely slow growing *Komagataella phaffii*"

### S1 | SUPPLEMENTARY METHODS

#### S1.1 | GSMM updates

The base genome-scale metabolic model (GSMM) used throughout this study was *i*MT1026v3, downloaded from BIOMODELS (Malik-Sheriff et al. 2020) `https://www.ebi.ac.uk/biomodels/MODEL1612130000`. Reactions were added to the model representing: the production of recombinant vHH, including cofactor requirements calculated according to (Stephanopoulos, Aristidou, & Nielsen 1998); a transport reaction for phosphatidylglycerol from the mitochondria to the cytoplasm for phospholipid composition (not present in the consensus biomass equation, but is in our quantification); and a sink reaction for glycogen to represent the consumption reaction. Additionally, the directionality of a number of reactions were altered. Reactions involving beta-glucose were blocked (GLUK, G6PI, G6PI3) as they are redundant to the reactions utilising alpha-glucose. All other biomass reactions present in the model (methanol and glycerol) were blocked. All other recombinant protein producing reactions present in the model were blocked.

#### S1.2 | Biomass composition

All data on biomass compositions and cultivation parameters were extracted from (Rebnegger et al. 2023). The data consisted of 12 sampling points: a triplicate chemostat cultivation at a set point of 0.1 $h^{-1}$ (C0.1), and a triplicate retentostat cultivation, with one sampling point during the initial chemostat phase (0.0), and 10 at later points during the retentostat phase. Biomass composition data was available for the 0.1 $h^{-1}$ chemostat cultivation, the chemostat phase before retentostat initiation and three other time points ~6, ~14 and ~28 days after initiation.

Compositional data was available for: total protein content; amino acid composition; glycogen, trehalose and total carbohydrate content; lipid composition; and, DNA and RNA content. Amino acid composition and lipid class composition were not available for the ~6 d sampling point, instead an average between the retentostat preceeding-chemostat and the ~14 4 values was assumed. For some biomass components we used a static composition based on the consensus glucose biomass equation (Tomàs-Gamisans et al. 2016). These included: the nucleotide compositions of DNA and RNA; the fatty acid chain composition used for the calculation of the lipid class composition; the composition of the different sphingolipid classes due to their different fatty acid compositions; and the composition of chitin-glucan complexes and their contribution to the total carbohydrate content.

The method used for lipid quantification defined a limit of quantification based on external multi-point standards and internal standardisation (Schoeny et al. 2021), which for lipids including ceramides and glucosylceramides led to both being below the limit of quantification for almost all sampling points. However, these lipids have been represented in the consensus glucose biomass composition (Tomàs-Gamisans et al. 2016) and we wanted to account for them in our compositions. From the lipid analysis data, we calculated the concentration of ceramides and glucosylceramides using the external multi-point standards without internal standardisation. We fitted a linear regression model to the area of the external standards, and observed good linearity. Using this model, we could calculate the concentration of ceramides and glucosylceramides for each sampling point. Due to the required alternative extraction, and the lack of a commercially available standard, the inositol containing lipids (IPC, MIPC and $M(IP)_2C$) were not quantified. As these were present in the consensus glucose composition, we chose to include them based on a linear relationship with glucosylceramide Tomàs-Gamisans et al. (2016). We assumed a 1:1.75:0.4:0.1 ratio of GlcCer:IPC:MIPC:M(IP)2C (unpublished data). Additionally, for the calculation of lipid composition, an average fatty acid was used in place of the R group, calculated from the reported fatty acid analysis previously used in *i*MT1026v3 (Grillitsch et al. 2014; Tomàs-Gamisans et al. 2016).

The amino acid composition was calculated, assuming that Glx was evenly split between glutamate and glutamine, whilst Asx was split evenly between aspartate and asparagine. Due to acid-hydrolysis affecting it's recovery, tryptophan was not quantified, so we assumed it's contribution to cell weight equal to tyrosine. A lysozyme standard was used as a reference % residue recovery in the amino acid analysis, we decided not to account for this correction in the calculation of the amino acid distribution.

We included many of the assumptions about molecular weights (MW) made in Tomàs-Gamisans et al. (2016) for calculating biomass compositions. The metabolite formula and mass were extracted from CheBI (Hastings et al. 2016) but to summarise:

(i) For the DNA composition, DNA nucleotides were represented as deoxynucleotide monophosphate residues

(ii) For the RNA composition, RNA nucleotides were represented as nucleotide monophosphate residues

(iii) Glycogen and D-glucan were considered as glucose residues without water. Mannan as 6-carbon mannose polymer and chitin as a single N-acetylglucosamine residue.

(iv) For the amino acid composition, amino acids were represented as residues.

Biomass compositions were calculated using the variable compositional data in combination with the static compositions described.

#### S1.3 | Exchange rate constraints

Oxygen uptake and carbon production rates were calculated through mass balancing of the gas entering and leaving the bioreactor. Using compositional data for the gas entering and leaving the system, as well as the different flow rates, oxygen uptake rate (OUR) and carbon production rate (CPR) were calculated according to (S1) and (S2), respectively.

$$OUR_{MB} = \frac{O_2in - O_2out}{V_L} = \frac{1}{V_L} \cdot \left( \frac{V.O_{2in}\% \cdot G_{in} - V.O_{2out}\% \cdot G_{out}}{V_m} \right) \quad \text{(S1)}$$

$$CPR_{MB} = \frac{CO_2out - CO_2in}{V_L} = \frac{1}{V_L} \cdot \left( \frac{V.CO_{2out}\% \cdot G_{out} - V.CO_{2in}\% \cdot G_{in}}{V_m} \right) \quad \text{(S2)}$$

Where $V_L$ volume of liquid in reactor, $G_{in}$ = gas flow rate in, $G_{out}$ = gas flow rate out and $V_m$ is the molar gas volume calculated by the ideal gas equation.

OUR and CPR for the chemostat cultivations were calculated at single time points once steady state was reached. For the retentostat cultivation, we observed cycling in the OUR and CPR. We smoothed these dynamics with a Gaussian process regressor such that one standard deviation of predicted value captured the cycling of the OUR and CPR. The kernel consisted of: a constant kernel, a radial basis function kernel and a white kernel. We observed that there was a large amount of instability in the OUR, which we assumed was due to an error with the oxygen detector and subsequently oxygen constraints were not used in this analysis.

Biomass-specific $CO_2$ exchange rates were calculated using the fitted function according to (S3).

$$q_{CO_2} = \frac{1}{C_X} \cdot \left( CPR_{MB} \cdot \frac{1000}{V_L} \right) \quad \text{(S3)}$$

The error associated with the biomass-specific exchange rates was propagated from the uncertainty of CPR via (S5), where $\delta q_{CO_2,random}$ is the sample standard deviation calculated from the three replicates, and here $n$ refers to the number of replicates, three.

$$\delta q_{CO_2,systematic} = \frac{1}{n} \cdot \sqrt{\sum_{i=1}^{n} \left( \frac{\delta CPR_{MB,i} \cdot \frac{1000}{V_L}}{C_{X,i}} \right)^2} \quad \text{(S4)}$$

$$\delta q_{CO_2,net} = \sqrt{(\delta q_{CO_2,random})^2 + (\delta q_{CO_2,systematic})^2} \quad \text{(S5)}$$

No uncertainty was quantified from the regression model, therefore the upper and lower bounds for glucose uptake, vHH production rate and growth rate were calculated from the average of the triplicate cultivations and the sample standard deviation.

### S2 | SUPPLEMENTARY RESULTS

#### S2.1 | Flux sampling convergence

Flux balance analysis (FBA) and its derivatives require the definition of an objective function, commonly the maximisation of biomass production. However, many solutions may satisfy the constraints and the steady state assumption. Flux variability analysis (FVA) has been used previously to assess the range of flux values (Torres et al. 2019); however, the minimal or maximal values returned from FVA do not necessarily lead to a feasible solution in combination with other minimal/maximal values. Whereas flux sampling is an unbiased approach that generates a chain of solutions, each of which satisfy the system constraints and provide additional information on the probability distributions of flux values (Herrmann et al. 2019).

To sample flux distributions, we first assessed the convergence of the optGp sampler (Megchelenbrink et al. 2014) using the cultivation data at 0.1 $h^{-1}$. We assessed sampler convergence for the 5 different biomass equations: the consensus and scaled consensus equations, and the respective 0.1 $h^{-1}$ biomass equation of the interpolated, fitted and derived methods. The upper bound of the model growth rate was set to 105 % of the regression model predicted growth rate, while the lower bound was set to 95 % of the lowest of either the maximal FBA growth rate or the regression model growth rate. Then, reactions that did not carry any flux were identified via FVA and removed from the model. 4 chains of 1,250 samples were stored, but a total of 5,000, 50,000, 500,000 or 5,000,000 samples were generated which represent thinning factors of 1, 10, 100 or 1,000, respectively. Then, solutions containing type III thermodynamically infeasible fluxes were converted to their nearest feasible solution (Desouki et al. 2015). The generated chains were assessed with a number of diagnostics to determine whether the sampler had converged, these were the Geweke diagnostic, rank normalized folded-split-$\hat{R}$ ($\hat{R}$) and

Effective sample size (ESS) (Herrmann et al. 2019; Vehtari et al. 2021).

Convergence of the sampler improved with more thinning. When thinning factors of 1 and 10 were used, there was a significant amount of autocorrelation and the sampler failed to converge. However, with thinning factors of 100 and 1,000, the majority of reactions converged (Table S8). We looked at a subset of reactions that we used in our analysis (glycolysis, pentose phosphate pathway (PPP), oxidative phosphorylation etc.) and observed that a thinning factor of 1,000 was necessary for all reactions, with all the biomass equation generation methods to converge. Additionally, for the same reactions we looked at a number of diagnostic plots including trace and rank plots, and plots of local ESS, quantile ESS and evolution ESS. A similar result was observed, that the sampler convergence was better at higher thinning factors. Sometimes we observed that at lower thinning factors, the sampler would stick in certain areas of the solution space and they would not be revisited by the sampler. However, regardless of the thinning factor, the majority of sampled fluxes occurred in the same range for regardless of the thinning factor. In our setup, the run-time increased by a maximum of 15 % between a thinning factor of 1 and 1,000. Based on this investigation, we chose to use a thinning factor of 1,000 for all later comparisons, in order to overcome the problems with autocorrelation and reach accurate conclusions about the metabolic flux distributions.

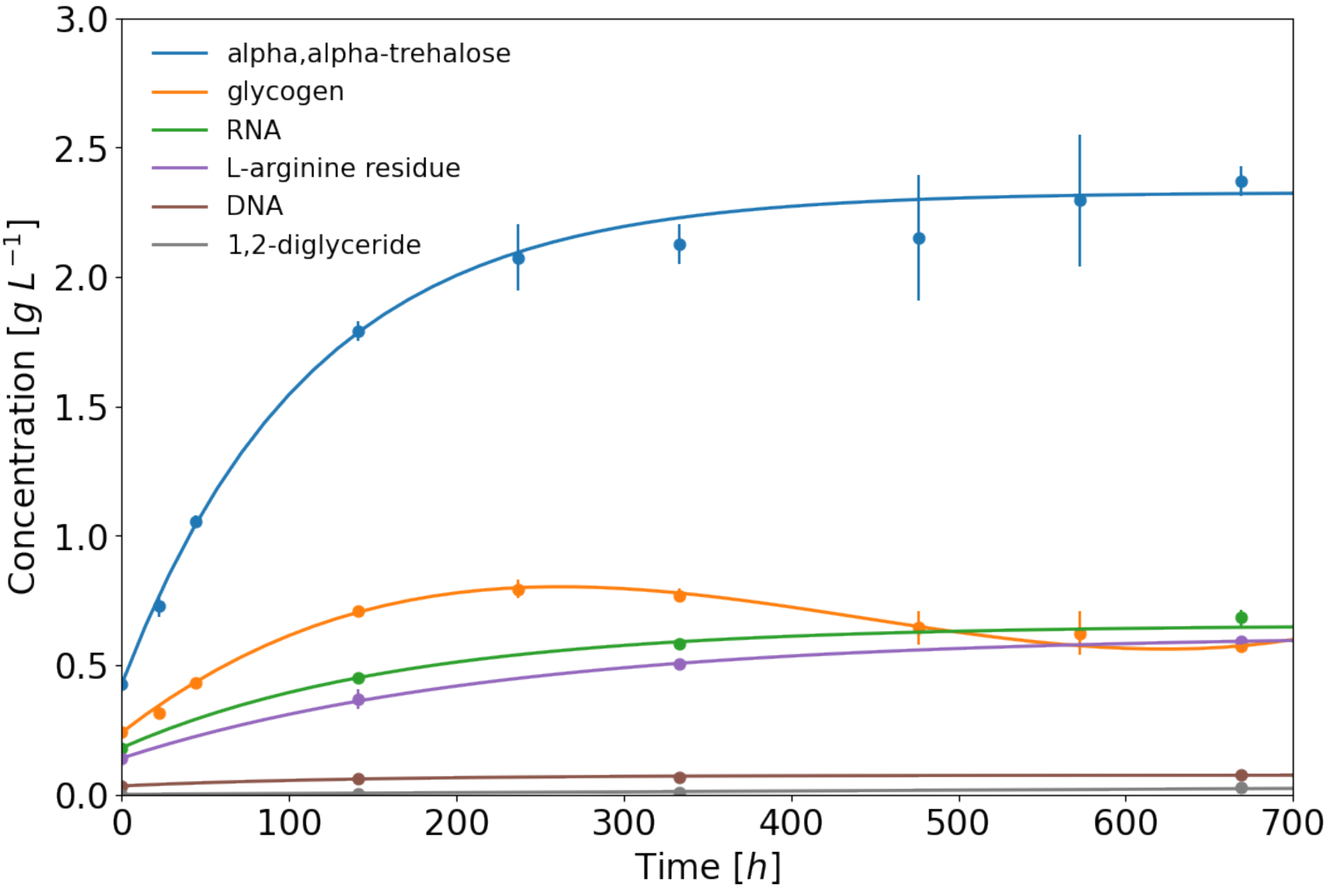


**FIGURE S1** Fitted functions to the total concentration of each biomass component as a function of time. Glycogen fitted with cubic polynomial, all others with an exponential asymptotic function (11). Points represent mean, vertical bars represent standard deviation. Shown are the most abundant components of the respective macromolecular classes; except for carbohydrates, where only the measured components are shown.

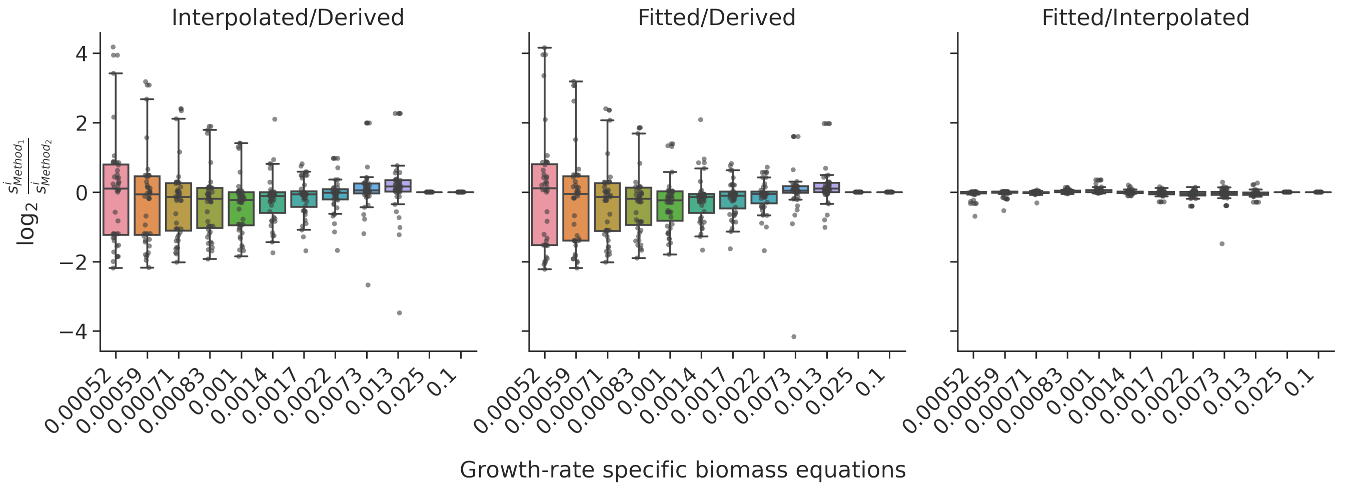


**FIGURE S2** $\text{Log}_2$ fold change in the stoichiometric coefficients between different biomass equation generation methods. In each comparison, the fold-change of each component was calculated at each growth rate between the two relevant growth-rate specific biomass equations.

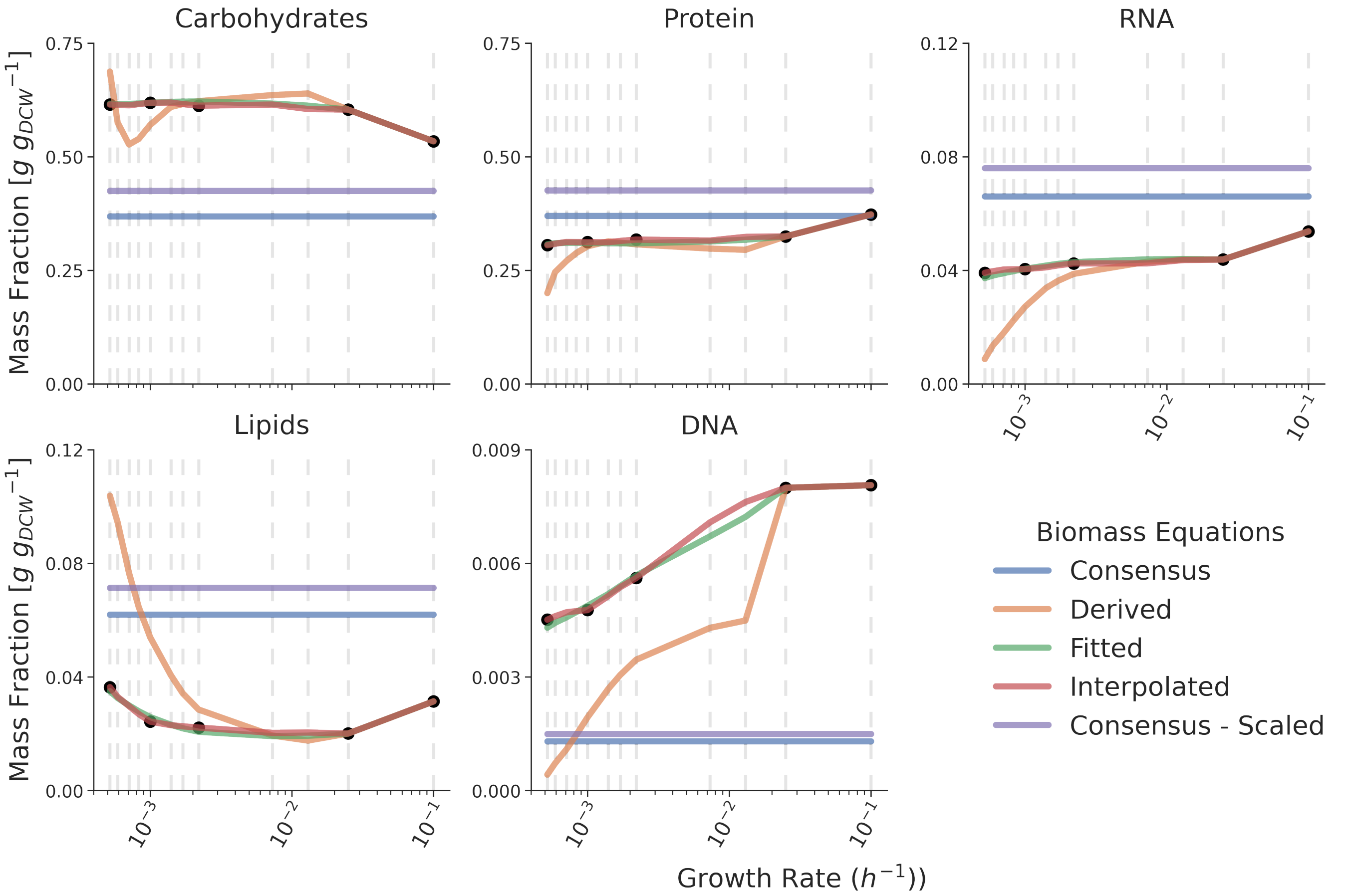


**FIGURE S3** The mass fraction of the five macromolecular classes across the investigated growth rates. The coloured lines represent the changing mass fractions of the biomass equations generated via the different methods, whereas the points represent the measured mass fraction. Dashed vertical lines represent the growth rates where growth rate specific biomass equations were generated.

**TABLE S1** Parameters used in regression model

| Parameter | Value | Description |
|---|---|---|
| $Y^{max}_{x/s}$ | 0.54 $g_{biomass}/g_{substrate}$ | Biomass-substrate yield |
| $Y^{max}_{p/s}$ | 0.609 $g_{product}/g_{substrate}$ | Product-substrate yield |
| v | $p_3(t; n_3)$ fit to each cultivation | Proportion of living cells |
| $C_{P,R}$ | $p_4(t; n_4)$ fit to each cultivation, $g\,L^{-1}$ | Product concentration in retentostat |
| $C_{S,R}$ | $0\,g\,L^{-1}$ | Substrate concentration in retentostat |
| $C_{S,MC}$ | $10\,g\,L^{-1}$ | Substrate concentration of chemostat media |
| $C_{S,MR}$ | $3.5\,g\,L^{-1}$ | Substrate concentration of retentostat media |
| $k_d$ | | Specific rate of inviable biomass generation |
| $D$ | $0.025\,h^{-1}$ | Dilution rate |
| $C_{X,v}/C_{X,d}$ | | Viable/Inviable biomass concentration |
| $C_X$ | | Total biomass concentration |
| $q_s$ | | Specific glucose uptake rate |
| $V_L$ | | Liquid volume in retentostat |
| $G_{in}$ | | Gas flow rate in |
| $G_{out}$ | | Gas flow rate out |

**TABLE S2** Statistics for adapted regression model fitting

| | $m_s$ [mg $g_{CDW}{}^{-1}$ $h^{-1}$] | Sum-of-squared Errors | $R^2$ | $R^2_{dead}$ | Growth Rate at end [$h^{-1}$] | Doubling time at end [d] | $q_P$ at end [mg $g_{CDW}{}^{-1}$ $h^{-1}$] |
|---|---|---|---|---|---|---|---|
| Cultivation 1 | 3.64 ± 7.59e-06 | 359 | 1 | 0.98 | 0.000529 | 54.5 | 0.0335 |
| Cultivation 2 | 3.76 ± 1.01e-05 | 852 | 0.999 | 0.901 | 0.000509 | 56.8 | 0.0364 |
| Cultivation 3 | 3.55 ± 8.38e-06 | 963 | 0.999 | 0.999 | 0.000502 | 57.5 | 0.0213 |

**TABLE S3** Measured Carbohydrate content [mg $g_{CDW}{}^{-1}$]

| | Sampling points | | | | | | | | | |
|---|---|---|---|---|---|---|---|---|---|---|
| Component | C0.1 | 0 | 1 | 2 | 6 | 10 | 14 | 20 | 24 | 28 |
| alpha,alpha-trehalose | 35.0 ± 0.48 | 89.0 ± 2.4 | 100.0 ± 6.1 | 130.0 ± 2.7 | 150.0 ± 3.1 | 150.0 ± 9.1 | 130.0 ± 4.8 | 120.0 ± 14.0 | 120.0 ± 13.0 | 120.0 ± 2.9 |
| glycogen | 54.0 ± 0.73 | 51.0 ± 1.4 | 45.0 ± 3.0 | 52.0 ± 2.7 | 58.0 ± 1.2 | 56.0 ± 2.6 | 48.0 ± 1.7 | 36.0 ± 3.7 | 33.0 ± 4.5 | 29.0 ± 0.71 |

**TABLE S4** Biomass compositions generated from experimental data and literature [mg $g_{CDW}^{-1}$]

| Component | C0.1 | 0 | 6 | 14 | 28 |
|---|---|---|---|---|---|
| | | | Sampling Point | | |
| (1->4)-beta-D-glucan | 290.0 ± 4.0 | 300.0 ± 8.0 | 260.0 ± 5.0 | 300.0 ± 10.0 | 320.0 ± 8.0 |
| (1->4)-beta-D-mannan | 62.0 ± 0.8 | 62.0 ± 2.0 | 62.0 ± 1.0 | 62.0 ± 2.0 | 62.0 ± 2.0 |
| 1,2-diglyceride | 0.39 ± 0.01 | 0.17 ± 0.01 | 0.41 ± 0.04 | 0.65 ± 0.05 | 1.3 ± 0.09 |
| 1-phosphatidyl-1D-myo-inositol(1-) | 2.1 ± 0.07 | 1.5 ± 0.1 | 1.8 ± 0.2 | 2.1 ± 0.2 | 2.9 ± 0.2 |
| alpha,alpha-trehalose | 35.0 ± 0.5 | 89.0 ± 2.0 | 150.0 ± 3.0 | 130.0 ± 5.0 | 120.0 ± 3.0 |
| Ceramide | 0.026 ± 0.0008 | 0.023 ± 0.002 | 0.027 ± 0.003 | 0.03 ± 0.002 | 0.093 ± 0.006 |
| chitin | 15.0 ± 0.2 | 15.0 ± 0.4 | 15.0 ± 0.3 | 15.0 ± 0.6 | 15.0 ± 0.4 |
| DNA | 6.9 ± 0.2 | 6.9 ± 0.4 | 4.9 ± 0.07 | 4.3 ± 0.1 | 4.0 ± 0.2 |
| ergosterol | 6.1 ± 0.2 | 4.2 ± 0.3 | 4.4 ± 0.4 | 4.5 ± 0.3 | 7.4 ± 0.5 |
| ergosteryl ester | 1.0 ± 0.03 | 0.27 ± 0.02 | 0.49 ± 0.05 | 0.72 ± 0.05 | 1.2 ± 0.08 |
| Glucosylceramide | 0.013 ± 0.0004 | 0.011 ± 0.0008 | 0.0079 ± 0.0008 | 0.0048 ± 0.0004 | 0.0094 ± 0.0006 |
| glycine residue | 11.0 ± 0.04 | 9.6 ± 0.2 | 9.5 ± 0.9 | 9.4 ± 0.2 | 9.3 ± 0.2 |
| glycogen | 54.0 ± 0.7 | 51.0 ± 1.0 | 58.0 ± 1.0 | 48.0 ± 2.0 | 29.0 ± 0.7 |
| IPC | 0.023 ± 0.0007 | 0.019 ± 0.001 | 0.014 ± 0.001 | 0.0084 ± 0.0006 | 0.017 ± 0.001 |
| L-alanine residue | 17.0 ± 0.07 | 14.0 ± 0.3 | 14.0 ± 1.0 | 14.0 ± 0.3 | 14.0 ± 0.3 |
| L-arginine residue | 29.0 ± 0.1 | 29.0 ± 0.6 | 30.0 ± 3.0 | 31.0 ± 0.7 | 30.0 ± 0.7 |
| L-asparagine residue | 16.0 ± 0.06 | 13.0 ± 0.3 | 14.0 ± 1.0 | 14.0 ± 0.3 | 14.0 ± 0.3 |
| L-aspartic acid residue | 16.0 ± 0.06 | 13.0 ± 0.3 | 14.0 ± 1.0 | 14.0 ± 0.3 | 15.0 ± 0.3 |
| L-cysteine residue | 3.7 ± 0.01 | 3.2 ± 0.06 | 3.3 ± 0.3 | 3.3 ± 0.07 | 3.2 ± 0.07 |
| L-glutamic acid residue | 30.0 ± 0.1 | 27.0 ± 0.5 | 26.0 ± 3.0 | 24.0 ± 0.5 | 20.0 ± 0.5 |
| L-glutamine residue | 30.0 ± 0.1 | 27.0 ± 0.5 | 25.0 ± 3.0 | 24.0 ± 0.5 | 20.0 ± 0.5 |
| L-histidine residue | 6.5 ± 0.03 | 6.6 ± 0.1 | 7.1 ± 0.7 | 7.5 ± 0.2 | 7.6 ± 0.2 |
| L-isoleucine residue | 12.0 ± 0.05 | 11.0 ± 0.2 | 11.0 ± 1.0 | 10.0 ± 0.2 | 10.0 ± 0.2 |
| L-leucine residue | 22.0 ± 0.09 | 19.0 ± 0.4 | 18.0 ± 2.0 | 18.0 ± 0.4 | 18.0 ± 0.4 |
| L-lysine residue | 22.0 ± 0.09 | 19.0 ± 0.4 | 20.0 ± 2.0 | 21.0 ± 0.5 | 21.0 ± 0.5 |
| L-methionine residue | 4.7 ± 0.02 | 4.4 ± 0.08 | 4.2 ± 0.4 | 4.1 ± 0.09 | 3.9 ± 0.09 |
| L-phenylalanine residue | 13.0 ± 0.05 | 11.0 ± 0.2 | 11.0 ± 1.0 | 11.0 ± 0.2 | 11.0 ± 0.2 |
| L-proline residue | 12.0 ± 0.05 | 11.0 ± 0.2 | 11.0 ± 1.0 | 11.0 ± 0.2 | 11.0 ± 0.2 |
| L-serine residue | 18.0 ± 0.07 | 15.0 ± 0.3 | 15.0 ± 1.0 | 14.0 ± 0.3 | 14.0 ± 0.3 |
| L-threonine residue | 17.0 ± 0.07 | 15.0 ± 0.3 | 15.0 ± 1.0 | 15.0 ± 0.3 | 15.0 ± 0.3 |
| L-tryptophan residue | 13.0 ± 0.05 | 11.0 ± 0.2 | 11.0 ± 1.0 | 11.0 ± 0.3 | 11.0 ± 0.2 |
| L-tyrosine residue | 11.0 ± 0.05 | 9.5 ± 0.2 | 9.7 ± 1.0 | 9.9 ± 0.2 | 9.4 ± 0.2 |
| L-valine residue | 14.0 ± 0.06 | 12.0 ± 0.2 | 12.0 ± 1.0 | 12.0 ± 0.3 | 12.0 ± 0.3 |
| lysophosphatidylcholine | 0.078 ± 0.002 | 0.062 ± 0.005 | 0.089 ± 0.009 | 0.12 ± 0.009 | 0.16 ± 0.01 |
| M(IP)2C | 0.0013 ± 4e-05 | 0.0011 ± 8e-05 | 0.00079 ± 8e-05 | 0.00048 ± 4e-05 | 0.00094 ± 6e-05 |
| MIPC | 0.0053 ± 0.0002 | 0.0044 ± 0.0003 | 0.0032 ± 0.0003 | 0.0019 ± 0.0001 | 0.0038 ± 0.0002 |
| phosphatidyl-L-serine(1-) | 0.49 ± 0.02 | 0.45 ± 0.03 | 0.51 ± 0.05 | 0.56 ± 0.04 | 0.73 ± 0.05 |
| phosphatidylcholine | 5.6 ± 0.2 | 4.5 ± 0.3 | 4.7 ± 0.5 | 5.0 ± 0.4 | 6.1 ± 0.4 |
| phosphatidylethanolamine | 9.0 ± 0.3 | 4.7 ± 0.3 | 5.4 ± 0.5 | 6.1 ± 0.5 | 8.1 ± 0.5 |
| phosphatidylglycerol | 0.77 ± 0.02 | 0.36 ± 0.03 | 0.38 ± 0.04 | 0.41 ± 0.03 | 0.63 ± 0.04 |
| RNA | 46.0 ± 0.5 | 38.0 ± 0.7 | 37.0 ± 0.6 | 36.0 ± 0.5 | 34.0 ± 2.0 |
| triglyceride | 1.1 ± 0.03 | 1.0 ± 0.08 | 1.2 ± 0.1 | 1.4 ± 0.1 | 3.0 ± 0.2 |
| zymosterol ester | 0.026 ± 0.0008 | 0.0 ± 0.0 | 0.064 ± 0.006 | 0.13 ± 0.01 | 0.24 ± 0.02 |

**TABLE S5** Growth rate specific biomass compositions [%]

| Macromolecule | Biomass Equation | Growth Rate $h^{-1}$ | | | | | | | | | | | |
|---|---|---|---|---|---|---|---|---|---|---|---|---|---|
| | | 0.1 | 0.025 | 0.013 | 0.0073 | 0.0022 | 0.0017 | 0.0014 | 0.001 | 0.00083 | 0.00071 | 0.00059 | 0.00052 |
| Carbohydrates | Consensus | | | | | | | 36.90 | | | | | |
| | Consensus - Scaled | | | | | | | 42.50 | | | | | |
| | Derived | 53.39 | 60.36 | 63.90 | 63.56 | 62.22 | 61.58 | 60.94 | 56.97 | 53.89 | 52.69 | 57.49 | 68.72 |
| | Fitted | 53.39 | 60.36 | 61.23 | 61.68 | 62.14 | 62.12 | 62.04 | 61.85 | 61.71 | 61.59 | 61.53 | 61.65 |
| | Interpolated | 53.39 | 60.36 | 60.48 | 61.47 | 61.22 | 61.57 | 61.90 | 61.86 | 61.58 | 61.29 | 61.44 | 61.47 |
| Protein | Consensus | | | | | | | 37.00 | | | | | |
| | Consensus - Scaled | | | | | | | 42.61 | | | | | |
| | Derived | 37.29 | 32.45 | 29.51 | 29.77 | 30.71 | 31.07 | 31.36 | 30.28 | 28.76 | 27.05 | 24.61 | 19.98 |
| | Fitted | 37.29 | 32.45 | 31.72 | 31.34 | 30.93 | 30.93 | 30.95 | 31.02 | 31.05 | 31.05 | 30.95 | 30.70 |
| | Interpolated | 37.29 | 32.45 | 32.35 | 31.55 | 31.76 | 31.46 | 31.18 | 31.19 | 31.22 | 31.24 | 30.84 | 30.52 |
| RNA | Consensus | | | | | | | 6.60 | | | | | |
| | Consensus - Scaled | | | | | | | 7.60 | | | | | |
| | Derived | 5.37 | 4.38 | 4.38 | 4.30 | 3.87 | 3.62 | 3.37 | 2.72 | 2.24 | 1.81 | 1.34 | 0.88 |
| | Fitted | 5.37 | 4.38 | 4.39 | 4.38 | 4.29 | 4.22 | 4.16 | 4.05 | 3.97 | 3.90 | 3.81 | 3.72 |
| | Interpolated | 5.37 | 4.38 | 4.36 | 4.24 | 4.24 | 4.17 | 4.10 | 4.04 | 4.04 | 4.03 | 3.96 | 3.91 |
| Lipids | Consensus | | | | | | | 6.20 | | | | | |
| | Consensus - Scaled | | | | | | | 7.14 | | | | | |
| | Derived | 3.14 | 2.01 | 1.76 | 1.94 | 2.85 | 3.42 | 4.06 | 5.40 | 6.47 | 7.66 | 9.44 | 10.38 |
| | Fitted | 3.14 | 2.01 | 1.93 | 1.92 | 2.07 | 2.19 | 2.32 | 2.59 | 2.79 | 3.00 | 3.26 | 3.50 |
| | Interpolated | 3.14 | 2.01 | 2.04 | 2.03 | 2.22 | 2.26 | 2.30 | 2.43 | 2.70 | 2.97 | 3.30 | 3.64 |
| DNA | Consensus | | | | | | | 0.13 | | | | | |
| | Consensus - Scaled | | | | | | | 0.15 | | | | | |
| | Derived | 0.81 | 0.80 | 0.45 | 0.43 | 0.35 | 0.31 | 0.27 | 0.19 | 0.15 | 0.11 | 0.07 | 0.04 |
| | Fitted | 0.81 | 0.80 | 0.72 | 0.67 | 0.57 | 0.54 | 0.52 | 0.49 | 0.47 | 0.46 | 0.44 | 0.43 |
| | Interpolated | 0.81 | 0.80 | 0.76 | 0.71 | 0.56 | 0.54 | 0.51 | 0.48 | 0.47 | 0.47 | 0.46 | 0.45 |

**TABLE S6** The 95 % confidence interval of sampled flux ratios relative to the glucose uptake rate. Negative numbers are wrapped with brackets.

| Equation | Growth Rate h$^{-1}$ | Reactions | | | | | | | | | | | | | | | | |
|---|---|---|---|---|---|---|---|---|---|---|---|---|---|---|---|---|---|---|
| | | G6PDH2 | PGI | PGMT | TRE6PS | PFK | PYK | PC | PDHa1 | CSm | ASPTA | ASPTAm | AKGMALtm | ASPGLU2m | MDH | MDHm | ME1m | Ex_nh4 |
| | 0.10000 | 0.44-0.47 | 0.34-0.37 | (0.19)-(0.19) | 0.00085-0.00085 | 0.58-0.59 | 1.1-1.2 | 0.37-0.39 | 0.66-0.69 | 0.53-0.56 | 0.0-0.65 | (0.94)-(0.8) | 0.96-1.1 | 0.8-0.94 | (1.0)-(0.9) | 1.4-1.5 | 0.0-0.033 | (0.61)-(0.6) |
| | 0.02500 | 0.37-0.47 | 0.35-0.46 | (0.17)-(0.17) | 0.00078-0.00079 | 0.6-0.63 | 1.2-1.2 | 0.23-0.36 | 0.75-0.79 | 0.63-0.67 | (0.08)-0.062 | (0.34)-(0.15) | 0.28-0.48 | 0.15-0.34 | (0.4)-(0.23) | 0.81-0.98 | 0.0-0.03 | (0.56)-(0.56) |
| | 0.01300 | 0.39-0.43 | 0.4-0.45 | (0.16)-(0.16) | 0.00072-0.00073 | 0.63-0.64 | 1.3-1.3 | 0.32-0.34 | 0.84-0.88 | 0.73-0.76 | 0.0-0.24 | (0.51)-(0.25) | 0.39-0.64 | 0.25-0.51 | (0.56)-(0.34) | 1.0-1.2 | 0.0-0.029 | (0.52)-(0.52) |
| | 0.00730 | 0.35-0.39 | 0.46-0.5 | (0.15)-(0.15) | 0.00067-0.00067 | 0.66-0.67 | 1.3-1.4 | 0.3-0.31 | 0.94-0.97 | 0.83-0.86 | 0.0-0.54 | (0.76)-(0.59) | 0.72-0.88 | 0.59-0.76 | (0.82)-(0.67) | 1.5-1.6 | 0.0-0.026 | (0.48)-(0.48) |
| | 0.00220 | 0.24-0.27 | 0.63-0.66 | (0.099)-(0.098) | 0.00044-0.00045 | 0.77-0.78 | 1.5-1.6 | 0.19-0.2 | 1.3-1.3 | 1.2-1.2 | 0.0-0.98 | (1.2)-(0.96) | 1.0-1.2 | 0.96-1.2 | (1.2)-(1.0) | 2.2-2.4 | 0.0-0.018 | (0.32)-(0.32) |
| Consensus - Scaled | 0.00170 | 0.2-0.23 | 0.69-0.71 | (0.086)-(0.085) | 0.00038-0.00039 | 0.8-0.81 | 1.6-1.6 | 0.17-0.18 | 1.4-1.4 | 1.3-1.3 | 0.0-1.1 | (1.3)-(1.1) | 1.2-1.4 | 1.1-1.3 | (1.4)-(1.1) | 2.4-2.7 | 0.0-0.015 | (0.28)-(0.28) |
| | 0.00140 | 0.18-0.21 | 0.72-0.74 | (0.076)-(0.075) | 0.00034-0.00034 | 0.82-0.83 | 1.7-1.7 | 0.15-0.16 | 1.5-1.5 | 1.4-1.4 | 0.0-1.2 | (1.3)-(1.2) | 1.3-1.4 | 1.2-1.3 | (1.3)-(1.2) | 2.6-2.7 | 0.0-0.014 | (0.25)-(0.25) |
| | 0.00100 | 0.15-0.17 | 0.77-0.79 | (0.053)-(0.053) | 0.00029-0.00029 | 0.86-0.87 | 1.7-1.7 | 0.13-0.13 | 1.6-1.6 | 1.5-1.5 | 0.0-1.3 | (1.4)-(1.3) | 1.4-1.5 | 1.3-1.4 | (1.5)-(1.4) | 2.9-2.9 | 0.0-0.012 | (0.21)-(0.21) |
| | 0.00083 | 0.14-0.16 | 0.8-0.82 | (0.04)-(0.04) | 0.00026-0.00026 | 0.88-0.89 | 1.8-1.8 | 0.11-0.12 | 1.6-1.6 | 1.6-1.6 | 0.0-1.4 | (1.5)-(1.4) | 1.5-1.6 | 1.4-1.5 | (1.5)-(1.4) | 3.0-3.1 | 0.0-0.01 | (0.19)-(0.19) |
| | 0.00071 | 0.13-0.29 | 0.68-0.84 | (0.032)-(0.032) | 0.00023-0.00024 | 0.85-0.9 | 1.8-1.8 | 0.053-0.11 | 1.6-1.7 | 1.6-1.6 | 1.4-1.8 | (1.8)-(1.5) | 1.5-1.9 | 1.5-1.8 | (1.9)-(1.5) | 3.1-3.4 | 0.0-0.0095 | (0.17)-(0.17) |
| | 0.00059 | 0.11-0.13 | 0.84-0.86 | (0.033)-(0.033) | 0.0002-0.0002 | 0.91-0.91 | 1.8-1.8 | 0.089-0.095 | 1.7-1.7 | 1.7-1.7 | 0.0-1.6 | (1.7)-(1.6) | 1.6-1.7 | 1.6-1.7 | (1.7)-(1.6) | 3.2-3.3 | 0.0-0.0083 | (0.15)-(0.15) |
| | 0.00052 | 0.1-0.11 | 0.84-0.86 | (0.041)-(0.04) | 0.00018-0.00019 | 0.9-0.91 | 1.8-1.8 | 0.081-0.088 | 1.7-1.7 | 1.7-1.7 | 1.5-1.7 | (1.7)-(1.6) | 1.6-1.8 | 1.6-1.7 | (1.8)-(1.6) | 3.2-3.4 | 0.0-0.0077 | (0.14)-(0.14) |
| | 0.10000 | 0.31-0.32 | 0.43-0.44 | (0.24)-(0.24) | 0.011-0.011 | 0.56-0.57 | 1.1-1.1 | 0.24-0.26 | 0.73-0.76 | 0.65-0.68 | 0.0-0.98 | (1.2)-(1.1) | 1.2-1.3 | 1.1-1.2 | (1.2)-(1.1) | 1.7-1.8 | 0.0-0.023 | (0.52)-(0.52) |
| | 0.02500 | 0.25-0.27 | 0.45-0.48 | (0.25)-(0.25) | 0.026-0.027 | 0.56-0.57 | 1.1-1.1 | 0.2-0.2 | 0.82-0.84 | 0.76-0.78 | 0.59-0.73 | (0.88)-(0.74) | 0.85-0.98 | 0.74-0.88 | (0.93)-(0.81) | 1.5-1.6 | 0.0-0.017 | (0.44)-(0.44) |
| | 0.01300 | 0.23-0.26 | 0.49-0.52 | (0.22)-(0.22) | 0.028-0.028 | 0.6-0.61 | 1.2-1.2 | 0.18-0.23 | 0.91-0.94 | 0.86-0.88 | 0.0-0.62 | (0.76)-(0.55) | 0.65-0.86 | 0.55-0.76 | (0.8)-(0.61) | 1.4-1.6 | 0.0-0.017 | (0.41)-(0.41) |
| | 0.00730 | 0.2-0.22 | 0.55-0.56 | (0.21)-(0.21) | 0.031-0.031 | 0.63-0.64 | 1.2-1.2 | 0.16-0.17 | 1.0-1.0 | 0.95-0.97 | 0.0-0.96 | (1.1)-(0.99) | 1.1-1.2 | 0.99-1.1 | (1.1)-(1.0) | 2.0-2.0 | 0.0-0.015 | (0.37)-(0.36) |
| | 0.00220 | 0.14-0.16 | 0.68-0.71 | (0.13)-(0.13) | 0.024-0.024 | 0.75-0.76 | 1.5-1.5 | 0.1-0.11 | 1.3-1.4 | 1.3-1.3 | 0.0-0.85 | (0.94)-(0.81) | 0.86-1.0 | 0.81-0.94 | (0.96)-(0.84) | 2.1-2.3 | 0.0-0.0099 | (0.24)-(0.24) |
| Interpolated | 0.00170 | 0.12-0.14 | 0.73-0.75 | (0.11)-(0.11) | 0.02-0.02 | 0.79-0.79 | 1.6-1.6 | 0.09-0.099 | 1.4-1.4 | 1.4-1.4 | 0.0-1.0 | (1.1)-(0.96) | 1.0-1.2 | 0.96-1.1 | (1.1)-(0.99) | 2.4-2.5 | 0.0-0.0085 | (0.21)-(0.21) |
| | 0.00140 | 0.11-0.13 | 0.75-0.77 | (0.1)-(0.1) | 0.018-0.018 | 0.81-0.82 | 1.6-1.6 | 0.08-0.087 | 1.5-1.5 | 1.5-1.5 | 0.0-1.1 | (1.2)-(1.1) | 1.1-1.2 | 1.1-1.2 | (1.2)-(1.1) | 2.6-2.7 | 0.0-0.0075 | (0.18)-(0.18) |
| | 0.00100 | 0.089-0.11 | 0.8-0.81 | (0.086)-(0.085) | 0.013-0.013 | 0.84-0.85 | 1.7-1.7 | 0.066-0.073 | 1.6-1.6 | 1.6-1.6 | 0.0-1.3 | (1.4)-(1.3) | 1.3-1.4 | 1.3-1.4 | (1.4)-(1.3) | 2.8-2.9 | 0.0-0.0061 | (0.15)-(0.15) |
| | 0.00083 | 0.079-0.092 | 0.82-0.84 | (0.075)-(0.074) | 0.011-0.011 | 0.86-0.87 | 1.7-1.7 | 0.06-0.065 | 1.6-1.6 | 1.6-1.6 | 0.0-1.4 | (1.4)-(1.3) | 1.4-1.5 | 1.3-1.4 | (1.4)-(1.4) | 3.0-3.0 | 0.0-0.0057 | (0.13)-(0.13) |
| | 0.00071 | 0.072-0.088 | 0.84-0.85 | (0.066)-(0.065) | 0.0093-0.0094 | 0.88-0.88 | 1.8-1.8 | 0.054-0.059 | 1.7-1.7 | 1.7-1.7 | 0.0-1.5 | (1.5)-(1.4) | 1.5-1.6 | 1.4-1.5 | (1.5)-(1.4) | 3.1-3.2 | 0.0-0.0052 | (0.12)-(0.12) |
| | 0.00059 | 0.066-0.077 | 0.86-0.87 | (0.058)-(0.058) | 0.0082-0.0083 | 0.89-0.9 | 1.8-1.8 | 0.049-0.053 | 1.7-1.7 | 1.7-1.7 | 0.0-1.5 | (1.6)-(1.5) | 1.5-1.6 | 1.5-1.6 | (1.6)-(1.5) | 3.2-3.3 | 0.0-0.0048 | (0.1)-(0.1) |
| | 0.00052 | 0.065-0.078 | 0.86-0.87 | (0.056)-(0.055) | 0.0077-0.0079 | 0.9-0.9 | 1.8-1.8 | 0.049-0.054 | 1.7-1.7 | 1.7-1.7 | 0.0-1.6 | (1.7)-(1.5) | 1.5-1.7 | 1.5-1.7 | (1.7)-(1.5) | 3.2-3.4 | 0.0-0.0051 | (0.11)-(0.1) |
| | 0.10000 | 0.31-0.32 | 0.43-0.44 | (0.24)-(0.24) | 0.011-0.011 | 0.56-0.57 | 1.1-1.1 | 0.24-0.26 | 0.74-0.76 | 0.65-0.67 | 0.0-0.96 | (1.1)-(1.1) | 1.2-1.3 | 1.1-1.1 | (1.2)-(1.1) | 1.7-1.8 | 0.0-0.023 | (0.52)-(0.52) |
| | 0.02500 | 0.25-0.27 | 0.46-0.48 | (0.25)-(0.25) | 0.026-0.027 | 0.57-0.57 | 1.1-1.1 | 0.19-0.21 | 0.82-0.84 | 0.76-0.78 | 0.0-0.74 | (0.88)-(0.75) | 0.85-0.99 | 0.75-0.88 | (0.93)-(0.81) | 1.5-1.6 | 0.0-0.017 | (0.44)-(0.44) |
| | 0.01300 | 0.22-0.24 | 0.5-0.52 | (0.22)-(0.22) | 0.033-0.033 | 0.6-0.61 | 1.2-1.2 | 0.18-0.19 | 0.92-0.94 | 0.87-0.88 | 0.46-0.56 | (0.68)-(0.59) | 0.68-0.78 | 0.59-0.68 | (0.73)-(0.64) | 1.5-1.6 | 0.0-0.015 | (0.4)-(0.4) |
| | 0.00730 | 0.2-0.21 | 0.55-0.56 | (0.2)-(0.2) | 0.034-0.034 | 0.63-0.64 | 1.2-1.2 | 0.16-0.17 | 1.0-1.0 | 0.96-0.98 | 0.0-0.97 | (1.1)-(0.97) | 1.1-1.2 | 0.97-1.1 | (1.1)-(1.0) | 1.9-2.1 | 0.0-0.014 | (0.37)-(0.37) |
| | 0.00220 | 0.13-0.16 | 0.69-0.71 | (0.13)-(0.13) | 0.024-0.024 | 0.75-0.76 | 1.5-1.5 | 0.11-0.11 | 1.3-1.4 | 1.3-1.3 | 0.0-0.85 | (0.96)-(0.8) | 0.86-1.0 | 0.8-0.96 | (0.98)-(0.83) | 2.1-2.3 | 0.0-0.0092 | (0.24)-(0.24) |
| Fitted | 0.00170 | 0.12-0.14 | 0.72-0.75 | (0.12)-(0.11) | 0.02-0.021 | 0.78-0.79 | 1.6-1.6 | 0.094-0.098 | 1.4-1.4 | 1.4-1.4 | 0.0-1.1 | (1.2)-(0.96) | 1.0-1.2 | 0.96-1.2 | (1.2)-(0.99) | 2.4-2.6 | 0.0-0.008 | (0.21)-(0.2) |
| | 0.00140 | 0.11-0.13 | 0.74-0.77 | (0.1)-(0.1) | 0.018-0.018 | 0.8-0.82 | 1.6-1.6 | 0.08-0.088 | 1.5-1.5 | 1.5-1.5 | 0.0-1.3 | (1.4)-(1.1) | 1.1-1.4 | 1.1-1.4 | (1.4)-(1.1) | 2.6-2.9 | 0.0-0.0073 | (0.19)-(0.18) |
| | 0.00100 | 0.089-0.1 | 0.8-0.81 | (0.085)-(0.084) | 0.014-0.014 | 0.84-0.85 | 1.7-1.7 | 0.07-0.073 | 1.6-1.6 | 1.6-1.6 | 0.0-1.3 | (1.4)-(1.3) | 1.3-1.4 | 1.3-1.4 | (1.4)-(1.3) | 2.8-2.9 | 0.0-0.0063 | (0.15)-(0.15) |
| | 0.00083 | 0.081-0.098 | 0.82-0.83 | (0.074)-(0.073) | 0.011-0.011 | 0.86-0.87 | 1.7-1.7 | 0.06-0.065 | 1.6-1.6 | 1.6-1.6 | 0.0-1.4 | (1.5)-(1.3) | 1.4-1.5 | 1.3-1.5 | (1.5)-(1.4) | 3.0-3.1 | 0.0-0.0057 | (0.13)-(0.13) |
| | 0.00071 | 0.072-0.085 | 0.84-0.85 | (0.066)-(0.065) | 0.0097-0.0099 | 0.88-0.88 | 1.8-1.8 | 0.054-0.059 | 1.7-1.7 | 1.7-1.7 | 0.0-1.5 | (1.6)-(1.4) | 1.4-1.6 | 1.4-1.6 | (1.6)-(1.4) | 3.1-3.2 | 0.0-0.0052 | (0.12)-(0.12) |
| | 0.00059 | 0.066-0.078 | 0.86-0.87 | (0.058)-(0.057) | 0.0083-0.0084 | 0.89-0.9 | 1.8-1.8 | 0.049-0.053 | 1.7-1.7 | 1.7-1.7 | 0.0-1.6 | (1.6)-(1.5) | 1.5-1.7 | 1.5-1.6 | (1.6)-(1.5) | 3.2-3.3 | 0.0-0.0049 | (0.1)-(0.1) |
| | 0.00052 | 0.066-0.17 | 0.77-0.87 | (0.056)-(0.055) | 0.0076-0.0077 | 0.86-0.9 | 1.8-1.8 | 0.031-0.053 | 1.7-1.7 | 1.7-1.7 | 0.0-1.8 | (1.8)-(1.5) | 1.5-1.8 | 1.5-1.8 | (1.8)-(1.5) | 3.2-3.5 | 0.0-0.005 | (0.1)-(0.1) |
| | 0.10000 | 0.31-0.32 | 0.43-0.44 | (0.24)-(0.24) | 0.011-0.011 | 0.56-0.57 | 1.1-1.1 | 0.24-0.26 | 0.73-0.76 | 0.65-0.67 | 0.0-0.96 | (1.1)-(1.1) | 1.2-1.3 | 1.1-1.1 | (1.2)-(1.1) | 1.7-1.8 | 0.0-0.023 | (0.52)-(0.52) |
| | 0.02500 | 0.24-0.28 | 0.44-0.48 | (0.25)-(0.25) | 0.026-0.027 | 0.56-0.58 | 1.1-1.1 | 0.19-0.21 | 0.82-0.85 | 0.76-0.78 | 0.0-1.0 | (1.1)-(0.63) | 0.74-1.3 | 0.63-1.1 | (1.2)-(0.69) | 1.4-1.9 | 0.0-0.017 | (0.45)-(0.44) |
| | 0.01300 | 0.21-0.24 | 0.49-0.52 | (0.22)-(0.21) | 0.058-0.059 | 0.59-0.6 | 1.1-1.2 | 0.17-0.18 | 0.92-0.94 | 0.87-0.89 | 0.0-0.54 | (0.66)-(0.51) | 0.61-0.76 | 0.51-0.66 | (0.71)-(0.57) | 1.4-1.5 | 0.0-0.013 | (0.38)-(0.38) |
| | 0.00730 | 0.19-0.21 | 0.54-0.56 | (0.2)-(0.2) | 0.049-0.049 | 0.63-0.63 | 1.2-1.2 | 0.15-0.17 | 1.0-1.0 | 0.96-0.98 | 0.0-0.84 | (0.96)-(0.84) | 0.93-1.0 | 0.84-0.96 | (1.0)-(0.89) | 1.8-1.9 | 0.0-0.012 | (0.35)-(0.35) |
| | 0.00220 | 0.14-0.28 | 0.56-0.7 | (0.14)-(0.13) | 0.022-0.022 | 0.71-0.76 | 1.5-1.5 | 0.11-0.12 | 1.3-1.3 | 1.3-1.3 | 0.0-1.4 | (1.5)-(0.79) | 0.85-1.5 | 0.79-1.5 | (1.5)-(0.82) | 2.1-2.7 | 0.0-0.01 | (0.24)-(0.23) |
| Derived | 0.00170 | 0.13-0.16 | 0.71-0.74 | (0.12)-(0.12) | 0.015-0.016 | 0.78-0.79 | 1.6-1.6 | 0.1-0.11 | 1.4-1.4 | 1.4-1.4 | 0.0-1.3 | (1.4)-(0.96) | 1.0-1.4 | 0.96-1.4 | (1.4)-(0.98) | 2.4-2.7 | 0.0-0.0096 | (0.21)-(0.2) |
| | 0.00140 | 0.12-0.38 | 0.5-0.76 | (0.11)-(0.11) | 0.011-0.011 | 0.73-0.81 | 1.5-1.6 | 0.053-0.099 | 1.4-1.5 | 1.4-1.5 | 0.0-1.6 | (1.6)-(1.1) | 1.1-1.7 | 1.1-1.6 | (1.6)-(1.1) | 2.5-3.0 | 0.0-0.0093 | (0.18)-(0.18) |
| | 0.00100 | 0.11-0.13 | 0.78-0.8 | (0.085)-(0.084) | 0.0056-0.0056 | 0.84-0.85 | 1.7-1.7 | 0.083-0.089 | 1.6-1.6 | 1.5-1.5 | 0.0-1.3 | (1.4)-(1.2) | 1.3-1.4 | 1.2-1.4 | (1.4)-(1.3) | 2.8-2.9 | 0.0-0.0091 | (0.15)-(0.14) |
| | 0.00083 | 0.11-0.12 | 0.81-0.82 | (0.072)-(0.072) | 0.0033-0.0034 | 0.87-0.87 | 1.7-1.7 | 0.077-0.083 | 1.6-1.6 | 1.6-1.6 | 0.0-1.4 | (1.4)-(1.3) | 1.4-1.5 | 1.3-1.4 | (1.5)-(1.4) | 2.9-3.0 | 0.0-0.0088 | (0.12)-(0.12) |
| | 0.00071 | 0.1-0.11 | 0.82-0.83 | (0.064)-(0.064) | 0.002-0.002 | 0.88-0.88 | 1.8-1.8 | 0.073-0.077 | 1.7-1.7 | 1.6-1.6 | 0.0-1.4 | (1.5)-(1.4) | 1.4-1.5 | 1.4-1.5 | (1.5)-(1.4) | 3.1-3.2 | 0.0-0.0086 | (0.1)-(0.1) |
| | 0.00059 | 0.095-0.11 | 0.83-0.84 | (0.065)-(0.063) | 0.00094-0.00095 | 0.89-0.89 | 1.8-1.8 | 0.066-0.07 | 1.7-1.7 | 1.7-1.7 | 1.4-1.6 | (1.7)-(1.5) | 1.5-1.7 | 1.5-1.7 | (1.7)-(1.5) | 3.2-3.3 | 0.0-0.0083 | (0.083)-(0.081) |
| | 0.00052 | 0.087-0.095 | 0.83-0.84 | (0.076)-(0.075) | 0.00042-0.00042 | 0.88-0.89 | 1.8-1.8 | 0.06-0.064 | 1.7-1.7 | 1.7-1.7 | 0.0-1.6 | (1.6)-(1.5) | 1.5-1.6 | 1.5-1.6 | (1.6)-(1.5) | 3.2-3.3 | 0.0-0.0079 | (0.068)-(0.065) |
| | 0.10000 | 0.43-0.44 | 0.37-0.39 | (0.18)-(0.18) | 0.00083-0.00083 | 0.59-0.6 | 1.2-1.2 | 0.36-0.38 | 0.7-0.72 | 0.57-0.6 | 0.0-1.1 | (1.4)-(1.3) | 1.5-1.6 | 1.3-1.4 | (1.5)-(1.4) | 1.9-2.0 | 0.0-0.032 | (0.59)-(0.59) |
| | 0.02500 | 0.4-0.42 | 0.41-0.43 | (0.17)-(0.17) | 0.00077-0.00078 | 0.62-0.62 | 1.2-1.3 | 0.33-0.36 | 0.78-0.8 | 0.66-0.68 | 0.0-0.97 | (1.2)-(1.1) | 1.3-1.4 | 1.1-1.2 | (1.3)-(1.2) | 1.8-1.9 | 0.0-0.03 | (0.55)-(0.55) |
| | 0.01300 | 0.37-0.39 | 0.45-0.47 | (0.16)-(0.16) | 0.00071-0.00071 | 0.65-0.65 | 1.3-1.3 | 0.31-0.33 | 0.87-0.9 | 0.76-0.79 | 0.0-0.99 | (1.2)-(1.1) | 1.2-1.4 | 1.1-1.2 | (1.3)-(1.2) | 1.9-2.0 | 0.0-0.028 | (0.51)-(0.51) |
| | 0.00730 | 0.34-0.35 | 0.5-0.52 | (0.14)-(0.14) | 0.00065-0.00065 | 0.68-0.68 | 1.3-1.4 | 0.28-0.3 | 0.97-0.99 | 0.87-0.89 | 0.0-1.4 | (1.6)-(1.5) | 1.6-1.7 | 1.5-1.6 | (1.6)-(1.6) | 2.4-2.4 | 0.0-0.025 | (0.47)-(0.47) |
| | 0.00220 | 0.22-0.24 | 0.67-0.69 | (0.09)-(0.09) | 0.00041-0.00041 | 0.79-0.8 | 1.6-1.6 | 0.18-0.19 | 1.3-1.4 | 1.3-1.3 | 0.0-0.98 | (1.1)-(1.0) | 1.1-1.2 | 1.0-1.1 | (1.2)-(1.0) | 2.3-2.4 | 0.0-0.016 | (0.3)-(0.29) |
| Consensus | 0.00170 | 0.18-0.2 | 0.72-0.74 | (0.077)-(0.077) | 0.00035-0.00035 | 0.82-0.83 | 1.6-1.7 | 0.16-0.16 | 1.4-1.5 | 1.4-1.4 | 0.0-1.1 | (1.2)-(1.1) | 1.2-1.3 | 1.1-1.2 | (1.3)-(1.2) | 2.5-2.6 | 0.0-0.014 | (0.25)-(0.25) |
| | 0.00140 | 0.17-0.19 | 0.75-0.76 | (0.068)-(0.068) | 0.00031-0.00031 | 0.84-0.85 | 1.7-1.7 | 0.13-0.14 | 1.5-1.5 | 1.5-1.5 | 0.0-1.3 | (1.4)-(1.3) | 1.3-1.4 | 1.3-1.4 | (1.4)-(1.3) | 2.7-2.8 | 0.0-0.012 | (0.22)-(0.22) |
| | 0.00100 | 0.14-0.15 | 0.8-0.82 | (0.046)-(0.045) | 0.00025-0.00026 | 0.88-0.89 | 1.8-1.8 | 0.11-0.12 | 1.6-1.6 | 1.6-1.6 | 0.0-1.4 | (1.5)-(1.4) | 1.4-1.6 | 1.4-1.5 | (1.5)-(1.4) | 3.0-3.1 | 0.0-0.01 | (0.19)-(0.18) |
| | 0.00083 | 0.12-0.14 | 0.83-0.85 | (0.032)-(0.032) | 0.00022-0.00023 | 0.9-0.91 | 1.8-1.8 | 0.098-0.1 | 1.7-1.7 | 1.6-1.6 | 0.0-1.5 | (1.6)-(1.5) | 1.5-1.7 | 1.5-1.6 | (1.6)-(1.5) | 3.1-3.3 | 0.0-0.0091 | (0.17)-(0.16) |
| | 0.00071 | 0.11-0.12 | 0.86-0.87 | (0.024)-(0.024) | 0.0002-0.0002 | 0.92-0.92 | 1.8-1.8 | 0.089-0.093 | 1.7-1.7 | 1.7-1.7 | 0.0-1.6 | (1.7)-(1.6) | 1.6-1.7 | 1.6-1.7 | (1.7)-(1.6) | 3.3-3.4 | 0.0-0.008 | (0.15)-(0.14) |
| | 0.00059 | 0.089-0.1 | 0.87-0.89 | (0.025)-(0.025) | 0.00016-0.00017 | 0.93-0.93 | 1.9-1.9 | 0.074-0.078 | 1.8-1.8 | 1.7-1.7 | 0.0-1.6 | (1.7)-(1.6) | 1.6-1.8 | 1.6-1.7 | (1.7)-(1.6) | 3.4-3.4 | 0.0-0.0068 | (0.12)-(0.12) |
| | 0.00052 | 0.081-0.091 | 0.88-0.89 | (0.033)-(0.032) | 0.00015-0.00015 | 0.92-0.93 | 1.8-1.9 | 0.066-0.072 | 1.8-1.8 | 1.7-1.7 | 0.0-1.7 | (1.8)-(1.6) | 1.7-1.8 | 1.6-1.8 | (1.8)-(1.7) | 3.4-3.5 | 0.0-0.0064 | (0.12)-(0.11) |

**TABLE S7** Major pathways of cofactor production. Shown are all pathways which contribute more than 1 % of the total supply across all growth rates.

| Cofactor | Equation | Subsystem | Growth Rate $h^{-1}$ 0.1 | 0.025 | 0.013 | 0.0073 | 0.0022 | 0.0017 | 0.0014 | 0.001 | 0.00083 | 0.00071 | 0.00059 | 0.00052 |
|---|---|---|---|---|---|---|---|---|---|---|---|---|---|---|
| atp_c | Consensus | Glycolysis/Gluconeogenesis | 19.2 | 18.5 | 17.4 | 16.1 | 14.2 | 13.7 | 13.4 | 13.0 | 12.8 | 12.7 | 12.5 | 12.5 |
| | | Transport, Mitochondrial | 80.8 | 81.5 | 82.6 | 83.9 | 85.8 | 86.3 | 86.6 | 87.0 | 87.2 | 87.3 | 87.5 | 87.5 |
| | Derived | Glycolysis/Gluconeogenesis | 17.4 | 16.2 | 15.4 | 14.7 | 13.5 | 13.3 | 13.0 | 12.9 | 12.7 | 12.6 | 12.5 | 12.4 |
| | | Transport, Mitochondrial | 82.6 | 83.8 | 84.6 | 85.3 | 86.4 | 86.7 | 87.0 | 87.1 | 87.3 | 87.4 | 87.5 | 87.6 |
| | Fitted | Glycolysis/Gluconeogenesis | 17.4 | 16.2 | 15.6 | 14.7 | 13.5 | 13.2 | 13.0 | 12.7 | 12.6 | 12.5 | 12.4 | 12.4 |
| | | Transport, Mitochondrial | 82.6 | 83.8 | 84.4 | 85.3 | 86.5 | 86.8 | 87.0 | 87.3 | 87.4 | 87.5 | 87.6 | 87.6 |
| | Interpolated | Glycolysis/Gluconeogenesis | 17.4 | 16.1 | 15.6 | 14.7 | 13.5 | 13.2 | 13.0 | 12.7 | 12.6 | 12.5 | 12.4 | 12.4 |
| | | Transport, Mitochondrial | 82.6 | 83.8 | 84.3 | 85.2 | 86.5 | 86.8 | 87.0 | 87.3 | 87.4 | 87.5 | 87.6 | 87.6 |
| | Consensus - Scaled | Glycolysis/Gluconeogenesis | 20.6 | 19.9 | 18.6 | 17.2 | 14.5 | 14.0 | 13.6 | 13.3 | 13.1 | 12.9 | 12.7 | 12.6 |
| | | Transport, Mitochondrial | 79.4 | 80.1 | 81.4 | 82.8 | 85.5 | 86.0 | 86.4 | 86.7 | 86.9 | 87.1 | 87.3 | 87.4 |
| atp_m | Consensus | Citric Acid Cycle | 4.3 | 4.9 | 5.3 | 5.3 | 6.3 | 6.4 | 6.4 | 6.5 | 6.5 | 6.5 | 6.5 | 6.6 |
| | | Oxidative Phosphorylation | 95.7 | 95.1 | 94.7 | 94.7 | 93.7 | 93.6 | 93.6 | 93.5 | 93.5 | 93.5 | 93.5 | 93.4 |
| | Derived | Citric Acid Cycle | 5.2 | 5.7 | 6.1 | 6.1 | 6.5 | 6.5 | 6.5 | 6.6 | 6.6 | 6.6 | 6.6 | 6.6 |
| | | Oxidative Phosphorylation | 94.8 | 94.3 | 93.9 | 93.9 | 93.5 | 93.5 | 93.5 | 93.4 | 93.4 | 93.4 | 93.4 | 93.4 |
| | Fitted | Citric Acid Cycle | 5.2 | 5.7 | 6.0 | 6.0 | 6.5 | 6.5 | 6.5 | 6.6 | 6.6 | 6.6 | 6.6 | 6.6 |
| | | Oxidative Phosphorylation | 94.8 | 94.3 | 94.0 | 94.0 | 93.5 | 93.5 | 93.5 | 93.4 | 93.4 | 93.4 | 93.4 | 93.4 |
| | Interpolated | Citric Acid Cycle | 5.2 | 5.7 | 6.0 | 6.0 | 6.5 | 6.5 | 6.5 | 6.6 | 6.6 | 6.6 | 6.6 | 6.6 |
| | | Oxidative Phosphorylation | 94.8 | 94.3 | 94.0 | 94.0 | 93.5 | 93.5 | 93.5 | 93.4 | 93.4 | 93.4 | 93.4 | 93.4 |
| | Consensus - Scaled | Citric Acid Cycle | 4.5 | 5.1 | 5.5 | 5.6 | 6.2 | 6.3 | 6.4 | 6.4 | 6.5 | 6.5 | 6.5 | 6.5 |
| | | Oxidative Phosphorylation | 95.5 | 94.9 | 94.5 | 94.4 | 93.8 | 93.7 | 93.6 | 93.6 | 93.5 | 93.5 | 93.5 | 93.5 |
| nadh_c | Consensus | Folate Metabolism | 0.0 | 0.0 | 0.0 | 0.0 | 0.0 | 0.0 | 0.5 | 0.4 | 0.0 | 0.0 | 0.0 | 0.0 |
| | | Glycine And Serine Metabolism | 5.3 | 3.8 | 3.5 | 4.0 | 1.9 | 1.6 | 1.8 | 1.1 | 1.0 | 0.9 | 0.7 | 0.7 |
| | | Glycolysis/Gluconeogenesis | 79.3 | 81.8 | 83.5 | 84.3 | 91.2 | 92.6 | 92.7 | 94.4 | 95.4 | 96.0 | 96.6 | 96.9 |
| | | Glyoxylate And Dicarboxylate Metabolism | 0.2 | 0.2 | 0.2 | 0.2 | 0.1 | 0.1 | 0.1 | 0.1 | 0.1 | 0.0 | 0.0 | 0.0 |
| | | Histidine Metabolism | 0.8 | 0.7 | 0.6 | 0.6 | 0.3 | 0.3 | 0.2 | 0.2 | 0.2 | 0.2 | 0.1 | 0.1 |
| | | Nucleotide Metabolism | 0.3 | 0.3 | 0.2 | 0.2 | 0.1 | 0.1 | 0.1 | 0.1 | 0.1 | 0.1 | 0.0 | 0.0 |
| | | Other Amino Acid Metabolism | 9.9 | 9.3 | 8.4 | 7.5 | 4.5 | 3.7 | 3.2 | 2.6 | 2.3 | 2.0 | 1.7 | 1.5 |
| | | Sterol Metabolism | 0.5 | 0.4 | 0.4 | 0.3 | 0.2 | 0.2 | 0.1 | 0.1 | 0.1 | 0.1 | 0.1 | 0.1 |
| | | Threonine And Lysine Metabolism | 1.3 | 1.2 | 1.1 | 1.0 | 0.6 | 0.5 | 0.4 | 0.3 | 0.3 | 0.3 | 0.2 | 0.2 |
| | | Valine, Leucine, And Isoleucine Metabolism | 2.5 | 2.3 | 2.1 | 1.9 | 1.1 | 0.9 | 0.8 | 0.7 | 0.6 | 0.5 | 0.4 | 0.4 |
| | Derived | Glycine And Serine Metabolism | 5.0 | 4.6 | 3.4 | 3.1 | 1.9 | 1.6 | 1.5 | 1.1 | 1.0 | 0.8 | 0.8 | 0.6 |
| | | Glycolysis/Gluconeogenesis | 84.5 | 87.1 | 89.8 | 90.6 | 93.7 | 94.4 | 94.7 | 95.3 | 95.6 | 95.9 | 96.2 | 96.5 |
| | | Glyoxylate And Dicarboxylate Metabolism | 0.1 | 0.1 | 0.1 | 0.1 | 0.0 | 0.1 | 0.1 | 0.1 | 0.1 | 0.1 | 0.1 | 0.1 |
| | | Histidine Metabolism | 0.7 | 0.7 | 0.7 | 0.6 | 0.4 | 0.3 | 0.3 | 0.3 | 0.2 | 0.2 | 0.2 | 0.1 |
| | | Nucleotide Metabolism | 0.3 | 0.2 | 0.2 | 0.2 | 0.1 | 0.1 | 0.1 | 0.0 | 0.0 | 0.0 | 0.0 | 0.0 |
| | | Other Amino Acid Metabolism | 5.2 | 3.8 | 3.2 | 3.0 | 2.3 | 2.1 | 2.1 | 2.1 | 2.1 | 2.1 | 2.0 | 2.0 |
| | | Sterol Metabolism | 0.3 | 0.2 | 0.1 | 0.1 | 0.1 | 0.1 | 0.1 | 0.1 | 0.1 | 0.2 | 0.2 | 0.2 |
| | | Threonine And Lysine Metabolism | 1.3 | 1.1 | 0.9 | 0.9 | 0.6 | 0.5 | 0.4 | 0.4 | 0.3 | 0.3 | 0.2 | 0.2 |
| | | Valine, Leucine, And Isoleucine Metabolism | 2.6 | 2.2 | 1.6 | 1.5 | 0.9 | 0.8 | 0.7 | 0.6 | 0.5 | 0.5 | 0.4 | 0.3 |
| | Fitted | Folate Metabolism | 0.0 | 0.0 | 0.0 | 0.0 | 0.0 | 0.0 | 0.0 | 0.0 | 0.0 | 0.0 | 0.2 | 0.0 |
| | | Glycine And Serine Metabolism | 5.0 | 4.4 | 3.7 | 3.2 | 1.8 | 1.6 | 1.5 | 1.1 | 1.0 | 0.8 | 0.8 | 0.9 |
| | | Glycolysis/Gluconeogenesis | 84.5 | 87.3 | 89.1 | 90.4 | 94.4 | 95.2 | 95.6 | 96.5 | 96.9 | 97.2 | 97.3 | 97.4 |
| | | Glyoxylate And Dicarboxylate Metabolism | 0.1 | 0.1 | 0.1 | 0.1 | 0.0 | 0.0 | 0.0 | 0.0 | 0.0 | 0.0 | 0.0 | 0.0 |
| | | Histidine Metabolism | 0.7 | 0.7 | 0.6 | 0.6 | 0.3 | 0.3 | 0.3 | 0.2 | 0.2 | 0.2 | 0.2 | 0.1 |
| | | Nucleotide Metabolism | 0.3 | 0.2 | 0.2 | 0.2 | 0.1 | 0.1 | 0.1 | 0.1 | 0.0 | 0.0 | 0.0 | 0.0 |
| | | Other Amino Acid Metabolism | 5.2 | 3.8 | 3.3 | 2.9 | 1.8 | 1.6 | 1.4 | 1.2 | 1.1 | 1.0 | 0.9 | 0.9 |
| | | Sterol Metabolism | 0.3 | 0.2 | 0.1 | 0.1 | 0.1 | 0.1 | 0.1 | 0.1 | 0.0 | 0.0 | 0.0 | 0.0 |
| | | Threonine And Lysine Metabolism | 1.3 | 1.1 | 1.0 | 0.9 | 0.5 | 0.4 | 0.4 | 0.3 | 0.3 | 0.2 | 0.2 | 0.2 |
| | | Valine, Leucine, And Isoleucine Metabolism | 2.6 | 2.2 | 1.9 | 1.6 | 0.9 | 0.8 | 0.7 | 0.6 | 0.5 | 0.4 | 0.4 | 0.4 |
| | Interpolated | Folate Metabolism | 0.0 | 1.0 | 0.0 | 0.0 | 0.0 | 0.0 | 0.0 | 0.0 | 0.0 | 0.0 | 0.0 | 0.0 |
| | | Glycine And Serine Metabolism | 5.0 | 4.6 | 3.8 | 3.3 | 1.9 | 1.6 | 1.4 | 1.3 | 1.0 | 0.9 | 0.8 | 0.8 |
| | | Glycolysis/Gluconeogenesis | 84.5 | 86.3 | 88.8 | 90.2 | 94.2 | 95.1 | 95.7 | 96.4 | 96.9 | 97.1 | 97.5 | 97.4 |
| | | Glyoxylate And Dicarboxylate Metabolism | 0.1 | 0.1 | 0.1 | 0.1 | 0.0 | 0.0 | 0.0 | 0.0 | 0.0 | 0.0 | 0.0 | 0.0 |
| | | Histidine Metabolism | 0.7 | 0.7 | 0.7 | 0.6 | 0.4 | 0.3 | 0.3 | 0.2 | 0.2 | 0.2 | 0.1 | 0.1 |
| | | Nucleotide Metabolism | 0.3 | 0.2 | 0.2 | 0.2 | 0.1 | 0.1 | 0.1 | 0.1 | 0.0 | 0.0 | 0.0 | 0.0 |
| | | Other Amino Acid Metabolism | 5.2 | 3.7 | 3.4 | 3.0 | 1.9 | 1.6 | 1.4 | 1.2 | 1.1 | 1.0 | 0.9 | 0.9 |
| | | Sterol Metabolism | 0.3 | 0.2 | 0.1 | 0.1 | 0.1 | 0.1 | 0.1 | 0.0 | 0.0 | 0.0 | 0.0 | 0.0 |
| | | Threonine And Lysine Metabolism | 1.3 | 1.1 | 1.0 | 0.9 | 0.5 | 0.4 | 0.4 | 0.3 | 0.3 | 0.2 | 0.2 | 0.2 |
| | | Valine, Leucine, And Isoleucine Metabolism | 2.6 | 2.2 | 2.0 | 1.7 | 1.0 | 0.8 | 0.7 | 0.6 | 0.5 | 0.4 | 0.4 | 0.4 |
| | Consensus - Scaled | Folate Metabolism | 0.0 | 0.0 | 0.0 | 0.0 | 0.0 | 0.0 | 0.0 | 0.5 | 0.0 | 0.0 | 0.4 | 0.0 |
| | | Glycine And Serine Metabolism | 4.2 | 5.0 | 3.5 | 3.7 | 2.0 | 1.7 | 1.5 | 1.3 | 1.1 | 1.2 | 0.9 | 1.1 |
| | | Glycolysis/Gluconeogenesis | 79.6 | 80.5 | 83.1 | 84.2 | 90.4 | 91.8 | 92.8 | 93.5 | 94.7 | 95.0 | 95.5 | 95.9 |
| | | Glyoxylate And Dicarboxylate Metabolism | 0.2 | 0.2 | 0.2 | 0.2 | 0.1 | 0.1 | 0.1 | 0.1 | 0.1 | 0.1 | 0.0 | 0.0 |
| | | Histidine Metabolism | 0.8 | 0.7 | 0.7 | 0.6 | 0.4 | 0.3 | 0.3 | 0.2 | 0.2 | 0.2 | 0.2 | 0.2 |
| | | Nucleotide Metabolism | 0.3 | 0.3 | 0.3 | 0.2 | 0.1 | 0.1 | 0.1 | 0.1 | 0.1 | 0.1 | 0.1 | 0.1 |
| | | Other Amino Acid Metabolism | 10.4 | 9.3 | 8.6 | 7.7 | 4.9 | 4.2 | 3.7 | 3.0 | 2.7 | 2.4 | 2.1 | 1.9 |
| | | Sterol Metabolism | 0.5 | 0.4 | 0.4 | 0.4 | 0.2 | 0.2 | 0.2 | 0.1 | 0.1 | 0.1 | 0.1 | 0.1 |
| | | Threonine And Lysine Metabolism | 1.4 | 1.2 | 1.1 | 1.0 | 0.6 | 0.5 | 0.5 | 0.4 | 0.4 | 0.3 | 0.3 | 0.3 |
| | | Valine, Leucine, And Isoleucine Metabolism | 2.6 | 2.3 | 2.2 | 1.9 | 1.2 | 1.0 | 0.9 | 0.8 | 0.7 | 0.6 | 0.5 | 0.5 |
| nadh_m | Consensus | Citric Acid Cycle | 68.1 | 64.2 | 62.6 | 65.0 | 58.1 | 58.5 | 58.9 | 59.1 | 59.3 | 59.3 | 59.5 | 59.6 |
| | | Glycine And Serine Metabolism | 0.6 | 0.0 | 0.0 | 0.3 | 0.1 | 0.0 | 0.1 | 0.0 | 0.0 | 0.0 | 0.0 | 0.0 |
| | | Glycolysis/Gluconeogenesis | 30.5 | 35.0 | 36.5 | 34.2 | 41.5 | 41.4 | 40.7 | 40.8 | 40.6 | 40.6 | 40.4 | 40.3 |
| | | Pyruvate Metabolism | 0.3 | 0.2 | 0.4 | 0.2 | 0.2 | 0.0 | 0.1 | 0.1 | 0.1 | 0.1 | 0.1 | 0.0 |
| | | Threonine And Lysine Metabolism | 0.6 | 0.5 | 0.4 | 0.3 | 0.2 | 0.1 | 0.1 | 0.1 | 0.1 | 0.1 | 0.1 | 0.0 |
| | Derived | Citric Acid Cycle | 64.1 | 59.7 | 57.1 | 59.2 | 57.1 | 57.5 | 58.2 | 58.3 | 58.5 | 58.8 | 59.0 | 59.1 |
| | | Glycine And Serine Metabolism | 0.0 | 0.2 | 0.0 | 0.0 | 0.0 | 0.0 | 0.1 | 0.0 | 0.0 | 0.0 | 0.0 | 0.0 |
| | | Glycolysis/Gluconeogenesis | 35.3 | 39.7 | 42.6 | 40.5 | 42.8 | 42.3 | 41.6 | 41.5 | 41.3 | 41.2 | 40.9 | 40.8 |
| | | Pyruvate Metabolism | 0.0 | 0.0 | 0.0 | 0.0 | 0.0 | 0.1 | 0.0 | 0.1 | 0.1 | 0.0 | 0.1 | 0.1 |
| | | Threonine And Lysine Metabolism | 0.5 | 0.4 | 0.3 | 0.3 | 0.2 | 0.1 | 0.1 | 0.1 | 0.1 | 0.1 | 0.0 | 0.0 |
| | Fitted | Citric Acid Cycle | 64.1 | 60.2 | 57.3 | 60.4 | 56.8 | 57.5 | 57.9 | 58.5 | 58.7 | 58.9 | 59.1 | 59.2 |
| | | Glycine And Serine Metabolism | 0.0 | 0.2 | 0.0 | 0.0 | 0.0 | 0.0 | 0.0 | 0.0 | 0.0 | 0.0 | 0.0 | 0.0 |
| | | Glycolysis/Gluconeogenesis | 35.3 | 39.2 | 42.0 | 39.3 | 43.0 | 42.3 | 41.9 | 41.4 | 41.2 | 41.0 | 40.8 | 40.7 |
| | | Pyruvate Metabolism | 0.0 | 0.0 | 0.3 | 0.0 | 0.1 | 0.1 | 0.1 | 0.1 | 0.0 | 0.0 | 0.0 | 0.0 |
| | | Threonine And Lysine Metabolism | 0.5 | 0.4 | 0.3 | 0.3 | 0.1 | 0.1 | 0.1 | 0.1 | 0.1 | 0.1 | 0.0 | 0.0 |
| | Interpolated | Citric Acid Cycle | 64.1 | 60.0 | 57.7 | 60.5 | 56.9 | 57.5 | 57.9 | 58.5 | 58.7 | 58.9 | 59.1 | 59.2 |
| | | Glycine And Serine Metabolism | 0.0 | 0.2 | 0.0 | 0.0 | 0.0 | 0.0 | 0.1 | 0.0 | 0.0 | 0.0 | 0.0 | 0.0 |
| | | Glycolysis/Gluconeogenesis | 35.1 | 39.1 | 41.6 | 39.3 | 42.9 | 42.4 | 41.9 | 41.4 | 41.3 | 41.0 | 40.8 | 40.7 |
| | | Pyruvate Metabolism | 0.3 | 0.2 | 0.3 | 0.0 | 0.1 | 0.0 | 0.1 | 0.0 | 0.0 | 0.0 | 0.0 | 0.0 |
| | | Threonine And Lysine Metabolism | 0.5 | 0.4 | 0.3 | 0.3 | 0.1 | 0.1 | 0.1 | 0.1 | 0.1 | 0.1 | 0.0 | 0.0 |
| | Consensus - Scaled | Citric Acid Cycle | 62.5 | 53.4 | 54.1 | 57.6 | 58.2 | 58.5 | 58.7 | 59.0 | 59.1 | 59.3 | 59.5 | 59.6 |
| | | Glycine And Serine Metabolism | 0.0 | 0.7 | 0.0 | 0.3 | 0.0 | 0.0 | 0.0 | 0.0 | 0.0 | 0.1 | 0.0 | 0.1 |
| | | Glycolysis/Gluconeogenesis | 36.0 | 44.5 | 44.8 | 41.3 | 41.4 | 41.2 | 41.0 | 40.8 | 40.7 | 40.5 | 40.4 | 40.2 |
| | | Pyruvate Metabolism | 0.7 | 0.7 | 0.6 | 0.4 | 0.2 | 0.2 | 0.1 | 0.1 | 0.1 | 0.1 | 0.1 | 0.1 |
| | | Threonine And Lysine Metabolism | 0.7 | 0.7 | 0.6 | 0.4 | 0.2 | 0.2 | 0.1 | 0.1 | 0.1 | 0.1 | 0.1 | 0.1 |
| nadph_c | Consensus | Folate Metabolism | 3.1 | 3.1 | 3.0 | 3.1 | 0.0 | 3.0 | 0.0 | 0.0 | 2.9 | 3.0 | 3.1 | 3.5 |
| | | Pentose Phosphate Pathway | 96.1 | 96.1 | 96.2 | 96.1 | 99.2 | 96.2 | 99.2 | 99.2 | 96.3 | 96.2 | 96.1 | 95.6 |
| | | Tyrosine, Tryptophan, And Phenylalanine Metabolism | 0.8 | 0.8 | 0.8 | 0.8 | 0.8 | 0.8 | 0.8 | 0.8 | 0.8 | 0.8 | 0.8 | 0.9 |
| | Derived | Folate Metabolism | 3.5 | 3.7 | 0.0 | 3.7 | 3.1 | 2.8 | 2.3 | 2.0 | 1.8 | 1.6 | 1.2 | 1.0 |
| | | Pentose Phosphate Pathway | 95.3 | 95.1 | 98.8 | 95.1 | 95.8 | 96.1 | 96.7 | 97.1 | 97.4 | 97.7 | 98.2 | 98.5 |
| | | Tyrosine, Tryptophan, And Phenylalanine Metabolism | 1.2 | 1.2 | 1.2 | 1.2 | 1.1 | 1.1 | 1.0 | 0.9 | 0.8 | 0.7 | 0.6 | 0.5 |
| | Fitted | Folate Metabolism | 3.6 | 3.6 | 3.7 | 3.6 | 3.4 | 3.3 | 3.2 | 3.0 | 3.0 | 3.0 | 0.0 | 3.1 |
| | | Pentose Phosphate Pathway | 95.3 | 95.2 | 95.1 | 95.2 | 95.5 | 95.6 | 95.7 | 95.9 | 96.0 | 95.9 | 98.9 | 95.8 |
| | | Tyrosine, Tryptophan, And Phenylalanine Metabolism | 1.2 | 1.1 | 1.2 | 1.2 | 1.1 | 1.1 | 1.1 | 1.1 | 1.1 | 1.1 | 1.1 | 1.1 |
| | Interpolated | Folate Metabolism | 3.5 | 0.0 | 3.3 | 3.5 | 3.3 | 3.3 | 3.2 | 3.1 | 3.2 | 3.0 | 2.9 | 3.1 |
| | | Glycerolipid Metabolism | 0.1 | 0.0 | 0.0 | 0.0 | 0.0 | 0.0 | 0.0 | 0.0 | 0.0 | 0.0 | 0.0 | 0.0 |
| | | Pentose Phosphate Pathway | 95.2 | 98.8 | 95.6 | 95.3 | 95.6 | 95.5 | 95.7 | 95.8 | 95.7 | 96.0 | 96.1 | 95.8 |
| | | Tyrosine, Tryptophan, And Phenylalanine Metabolism | 1.2 | 1.2 | 1.1 | 1.2 | 1.1 | 1.1 | 1.1 | 1.1 | 1.1 | 1.1 | 1.1 | 1.1 |
| | Consensus - Scaled | Folate Metabolism | 3.0 | 2.9 | 2.8 | 2.8 | 2.9 | 2.9 | 2.9 | 0.0 | 2.9 | 0.0 | 0.0 | 3.3 |
| | | Pentose Phosphate Pathway | 96.2 | 96.3 | 96.4 | 96.4 | 96.3 | 96.3 | 96.3 | 99.2 | 96.3 | 99.2 | 99.2 | 95.8 |
| | | Tyrosine, Tryptophan, And Phenylalanine Metabolism | 0.8 | 0.8 | 0.8 | 0.8 | 0.8 | 0.8 | 0.8 | 0.8 | 0.8 | 0.8 | 0.8 | 0.8 |
| nadph_m | Consensus | Citric Acid Cycle | 0.0 | 95.1 | 95.4 | 0.0 | 47.3 | 70.1 | 0.0 | 98.9 | 98.6 | 100.0 | 100.0 | 100.0 |
| | | Folate Metabolism | 100.0 | 4.9 | 4.6 | 100.0 | 52.7 | 2.3 | 100.0 | 1.1 | 1.4 | 0.0 | 0.0 | 0.0 |
| | | Pyruvate Metabolism | 0.0 | 0.0 | 0.0 | 0.0 | 0.0 | 27.6 | 0.0 | 0.0 | 0.0 | 0.0 | 0.0 | 0.0 |
| | Derived | Citric Acid Cycle | 95.2 | 56.6 | 54.2 | 86.3 | 93.8 | 94.7 | 0.0 | 93.3 | 88.9 | 52.9 | 0.0 | 20.8 |
| | | Folate Metabolism | 4.8 | 43.4 | 29.7 | 13.7 | 6.2 | 5.3 | 100.0 | 6.7 | 11.1 | 9.5 | 100.0 | 79.2 |
| | | Pyruvate Metabolism | 0.0 | 0.0 | 16.1 | 0.0 | 0.0 | 0.0 | 0.0 | 0.0 | 0.0 | 37.6 | 0.0 | 0.0 |
| | Fitted | Citric Acid Cycle | 95.0 | 56.6 | 92.0 | 73.0 | 96.3 | 92.8 | 52.8 | 53.5 | 97.3 | 95.9 | 61.8 | 9.1 |
| | | Folate Metabolism | 5.0 | 43.4 | 8.0 | 1.6 | 3.7 | 7.2 | 47.2 | 46.5 | 2.7 | 4.1 | 38.2 | 90.9 |
| | | Pyruvate Metabolism | 0.0 | 0.0 | 0.0 | 25.4 | 0.0 | 0.0 | 0.0 | 0.0 | 0.0 | 0.0 | 0.0 | 0.0 |
| | Interpolated | Citric Acid Cycle | 95.1 | 56.6 | 73.1 | 73.7 | 60.5 | 95.0 | 46.3 | 24.1 | 72.5 | 28.4 | 32.6 | 100.0 |
| | | Folate Metabolism | 4.9 | 43.4 | 26.9 | 1.1 | 39.5 | 5.0 | 53.7 | 75.9 | 3.1 | 71.6 | 67.4 | 0.0 |
| | | Pyruvate Metabolism | 0.0 | 0.0 | 0.0 | 25.2 | 0.0 | 0.0 | 0.0 | 0.0 | 24.4 | 0.0 | 0.0 | 0.0 |
| | Consensus - Scaled | Citric Acid Cycle | 94.7 | 0.0 | 97.6 | 0.0 | 96.3 | 96.3 | 96.6 | 98.6 | 98.0 | 0.0 | 100.0 | 0.0 |
| | | Folate Metabolism | 5.3 | 100.0 | 2.4 | 100.0 | 3.7 | 3.7 | 3.4 | 1.4 | 2.0 | 100.0 | 0.0 | 100.0 |

**TABLE S8** Diagnostics summary of flux sampling thinning

| | | Total no. of fails across 4 chains | | | % of reactions failed | | |
|---|---|---|---|---|---|---|---|
| Thinning Factor | Biomass Composition | Geweke | $\hat{R}$ | Bulk ESS | Geweke | $\hat{R}$ | Bulk ESS |
| 1 | Consensus | 828 | 460 | 745 | 93.6 | 52.0 | 84.2 |
| | Derived | 861 | 689 | 671 | 96.4 | 77.2 | 75.1 |
| | Fitted | 876 | 210 | 758 | 96.1 | 23.0 | 83.1 |
| | Interpolated | 888 | 300 | 683 | 93.2 | 31.5 | 71.7 |
| | Consensus - Scaled | 828 | 685 | 672 | 97.4 | 80.6 | 79.1 |
| 10 | Consensus | 842 | 280 | 518 | 94.3 | 31.4 | 58.0 |
| | Derived | 915 | 272 | 590 | 90.2 | 26.8 | 58.2 |
| | Fitted | 928 | 416 | 599 | 94.6 | 42.4 | 61.1 |
| | Interpolated | 888 | 584 | 665 | 93.4 | 61.4 | 69.9 |
| | Consensus - Scaled | 836 | 512 | 575 | 93.3 | 57.1 | 64.2 |
| 100 | Consensus | 711 | 50 | 21 | 73.4 | 5.2 | 2.2 |
| | Derived | 511 | 15 | 33 | 57.2 | 1.7 | 3.7 |
| | Fitted | 645 | 10 | 8 | 70.0 | 1.1 | 0.9 |
| | Interpolated | 602 | 44 | 6 | 65.1 | 4.8 | 0.6 |
| | Consensus - Scaled | 825 | 80 | 18 | 84.8 | 8.2 | 1.8 |
| 1000 | Consensus | 302 | 2 | 0 | 34.3 | 0.2 | 0.0 |
| | Derived | 355 | 28 | 5 | 38.3 | 3.0 | 0.5 |
| | Fitted | 426 | 56 | 19 | 46.5 | 6.1 | 2.1 |
| | Interpolated | 346 | 4 | 2 | 38.3 | 0.4 | 0.2 |
| | Consensus - Scaled | 348 | 0 | 0 | 37.1 | 0.0 | 0.0 |
